# NSUN2-mediated m^5^C methylation of IRF3 mRNA negatively regulates type I interferon responses

**DOI:** 10.1101/2021.07.09.451748

**Authors:** Hongyun Wang, Lu Zhang, Cong Zeng, Jiangpeng Feng, Dehe Wang, Miao He, Ao Jiang, Yuanyuan Cao, Xiao Guo, Jiejie Liu, Kun Yan, Hao Tang, Ke Xu, Chengpeng Fan, Deyin Guo, Ke Lan, Yu Zhou, Yu Chen

## Abstract

5-Methylcytosine (m^5^C) is a widespread post-transcriptional RNA modification and is reported to be involved in manifold cellular responses and biological processes through regulating RNA metabolism. However, its regulatory role in antiviral innate immunity has not yet been elucidated. Here, we report that NSUN2, a typical m^5^C methyltransferase, can negatively regulate type I interferon responses during viral infection. NSUN2 specifically mediates m^5^C methylation of *IRF3* mRNA and accelerates its degradation, resulting in low levels of IRF3 and downstream IFN-β production. Knockout or knockdown of NSUN2 could enhance type I interferon responses and downstream ISG expression after viral infection *in vitro*. And *in vivo*, the antiviral innate responses is more dramatically enhanced in *Nsun2^+/−^* mice than in *Nsun2^+/+^* mice. Four highly m^5^C methylated cytosines in *IRF3* mRNA were identified, and their mutation could enhance the cellular *IRF3* mRNA levels. Moreover, infection with Sendai virus (SeV), vesicular stomatitis virus (VSV), herpes simplex virus 1 (HSV-1), Zika virus (ZIKV), or especially SARS-CoV-2 resulted in a reduction in endogenous levels of NSUN2. Together, our findings reveal that NSUN2 serves as a negative regulator of interferon response by accelerating the fast turnover of *IRF3* mRNA, while endogenous NSUN2 levels decrease after viral infection to boost antiviral responses for the effective elimination of viruses. Our results suggest a paradigm of innate antiviral immune responses ingeniously involving NSUN2-mediated m^5^C modification.

## Introduction

RNA modification is an important post-transcriptional modification process. To date, more than 100 types of chemical modifications to various types of RNAs have been recorded (1). Among these RNA modifications, N6-methyladenosine (m^6^A) and 5-methylcytosine (m^5^C) are ubiquitous, and have led to an increasing appreciation that RNA methylation can functionally regulate gene expression and cellular activity (2-4). The methyltransferase (writer), demethylase (eraser), and effector (reader) play coordinating roles in RNA metabolism, such as splicing, degradation, and translation (5-9). Recently, it was found that m^6^A methylation could negatively regulate interferon response by inducing *IFNB* mRNA degradation (10, 11). It was reported that m^6^A RNA-modification-mediated downregulation of the OGDH-itaconate pathway reprograms cellular metabolism to inhibit viral replication (12). Another study demonstrated that ALKBH5, an m^6^A demethylase, could be recruited by DDX46 and then erase the m^6^A modification in *MAVS*, *TRAF3*, and *TRAF6* transcripts, thereby enforcing their retention in the nucleus and leading to their decreased translation, resulting in inhibited type I interferon production (13). Additionally, nuclear hnRNPA2B1 facilitates m^6^A modification and nucleocytoplasmic trafficking of *CGAS*, *IFI16*, and *STING* mRNAs, resulting in amplification of the innate immune response to DNA viruses (14). At present, m^5^C is not well studied compared to m^6^A. The primary writers for m^5^C methylation of RNA in animals have been proposed to be NSUN2 and TRDMT1 (DNMT2) (15, 16). NSUN2 is reported to regulate the expression of numerous genes by methylating their mRNAs and thereby affecting their degradation or translation (17-20). Another report emphasized the transcriptome-wide role of NSUN2 as a major methyltransferase of the m^5^C epitranscriptomic mark and presented compelling evidence for the functional interdependence of mRNA m^5^C methylation and mRNA translation (21). Furthermore, it was reported that ALYREF and YBX1 served as potential m^5^C readers that could recognize m^5^C-modified mRNA and mediate mRNA export from the nucleus or affect the stability of their target mRNAs (22-26). Nevertheless, the demethylases responsible for removing m^5^C methylation on RNA have not yet been identified. Moreover, whether the m^5^C modification participates in the regulation of antiviral innate immunity, similarly to m^6^A modification, and especially in regulating the production of type I interferon responses, remains to be defined.

Elicitation of type I interferons (IFNs) by viruses or other pathogens plays an extremely critical role in innate immunity. The induction of type I interferons is primarily controlled at the level of gene transcription, wherein the interferon regulatory factor (IRF) family of transcription factors plays a central role (27-30). Interferon regulatory factor 3 (IRF3) acts as a master transcription factor responsible for the induction of type I interferons and is essential for the establishment of antiviral innate immunity (31, 32). After viral infection, IRF3 is phosphorylated by the kinases TBK1 and IKKε on its C-terminal and undergoes a conformational change and homodimerization, which leads to its translocation to the nucleus and subsequent association with the interferon-stimulated response elements of target genes (33, 34). Because of its pivotal role in the induction of type I interferons, the transcription factor IRF3 requires sophisticated regulation in order to effectively maintain immune homeostasis after viral infection. It has been reported that a great deal of regulators of IRF3 participate in maintaining the appropriate amounts of type I interferons stimulated by viral infection (35-38). The reported regulators of IRF3 mostly induce changes in the phosphorylation levels or quantity of IRF3 protein, which then affects type I interferon responses and downstream ISG. Most reports mainly focus on the regulation of IRF3 at the protein level. However, there are few reports about the regulation of IRF3 at the mRNA level, especially involving epigenetic modification.

Herein, we revealed that NSUN2, a typical RNA m^5^C methyltransferase, serves as a negative regulator of type I interferon responses in antiviral innate immunity. We found that NSUN2 could specifically mediate m^5^C methylation of *IRF3* mRNA and accelerate its degradation, and that knockout or knockdown of NSUN2 could elevate both mRNA and protein levels of IRF3 and thus amplify type I interferon responses and downstream ISG expression after viral infection. Four highly m^5^C-methylated cytosines in *IRF3* mRNA were identified using bisulfite RNA sequencing, and the mutation of these cytosines could enhance the IRF3 levels and IFN-β production. We outline a paradigm of innate immune responses to viral infection in which genes are ingeniously regulated by epigenetic modification.

## Results

### NSUN2 negatively regulates type I interferon responses

To explore the function of RNA methyltransferase or demethylases involved in type I interferon responses, we knocked down different RNA methyltransferases or demethylases in HEK293T cells using small interfering RNAs (siRNAs) and detected the endogenous levels of *IFNB* mRNA. We found that compared with other methyltransferases or demethylases, knockdown of NSUN2 could more dramatically enhance endogenous *IFNB* mRNA levels (**Fig. 1a**). To confirm the impact of NSUN2 on type I interferon responses, we examined the effect of exogenous NSUN2 expression and found that it could inhibit the activation of IFN-β promoter activity induced by Sendai virus (SeV) in a dose-dependent manner (**Fig. 1b**). Exogenous NSUN2 expression could also inhibit the activation of IFN-β promoter activity induced by different stimulants (**Fig. 1c**). In NSUN2 knockdown HEK293T cells, the SeV-induced increase in endogenous *IFNB* mRNA levels was dramatically enhanced as was the mRNA levels of downstream *ISG15* and *CXCL10* (**Fig. 1d**). Moreover, SeV-induced type I interferon responses were significantly enhanced in NSUN2 knockout HEK293T cells (**Fig. 1e**) and A549 cells (**Supplementary Fig S1a**).

**Figure 1.**
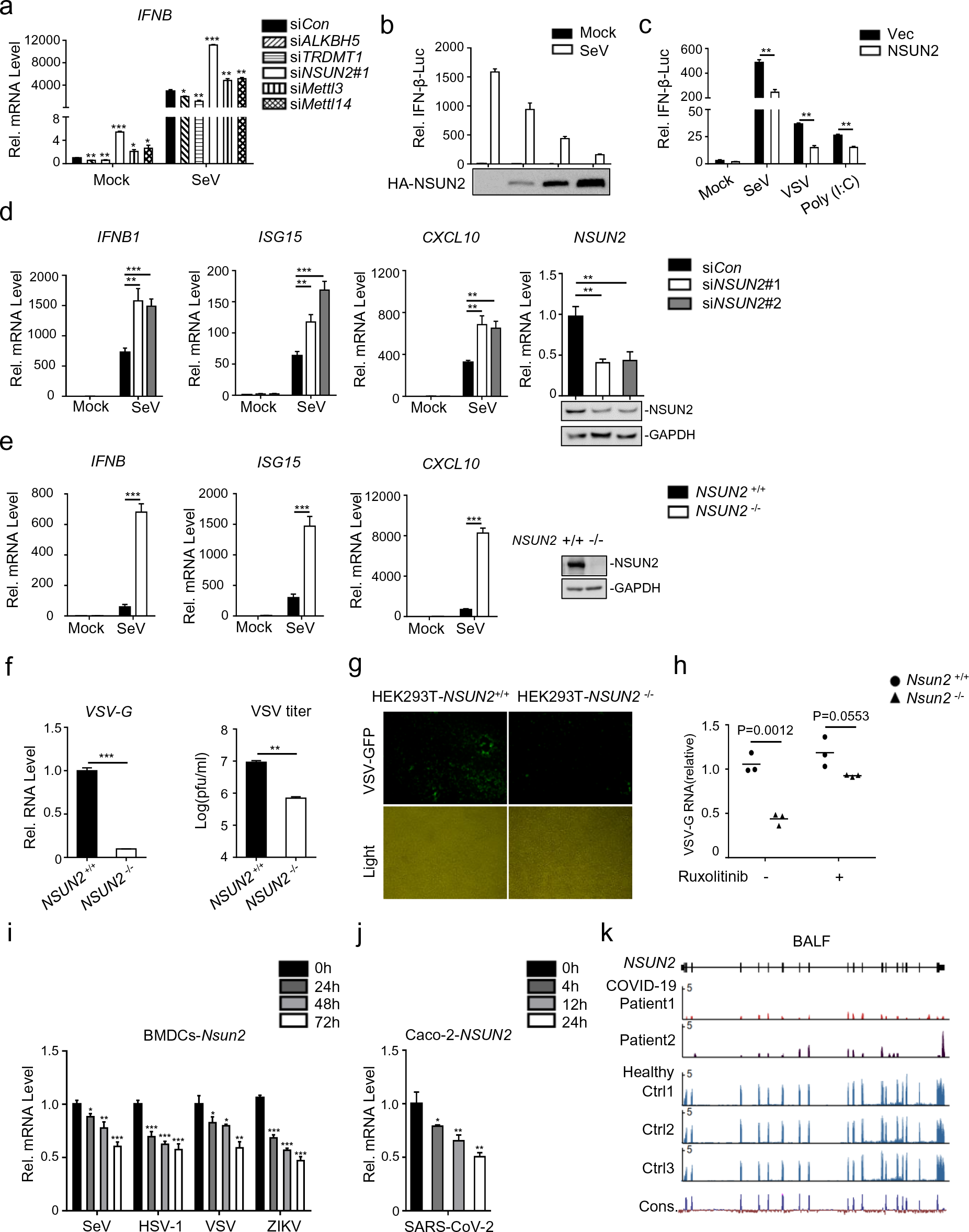
NSUN2 negatively regulates antiviral innate type I interferon responses. (a) qPCR analysis of *IFNB* mRNA in HEK293T cells transfected with siControl or siRNAs targeting different RNA methyltransferases or demethylases for 36 h, with or without infection by SeV for another 8 h. (b) Dual-luciferase assay analyzing IFN-β promoter activity (IFN-β-Luc) in HEK293T cells in 24-well plates transfected for 24 h with 100 ng IFN-β-Luc plasmid and 20 ng *Renilla* luciferase plasmid (RL-TK) along with vector or increasing amounts (0, 0.1, 0.2, and 0.5 μg) of plasmid encoding NSUN2, with or without infection by SeV, for another 10 h. (c) Dual-luciferase analysis of IFN-β-Luc in HEK293T cells in 24-well plates transfected for 24 h with vector (Vec) or NSUN2, with or without infection by SeV or VSV for another 10 h, or transfected with poly (I:C) (1 µg/mL) for another 10 h. (d) qPCR analysis of *IFNB*, *ISG15*, *CXCL10* and *NSUN2* mRNA in HEK293T cells transfected with siControl or siRNAs targeting NSUN2, with or without infection by SeV for 8 h. Immunoblot analysis shows knockdown efficiency of siRNAs targeting NSUN2. (e) qPCR analysis of *IFNB*, *ISG15* and *CXCL10* mRNA in wild-type HEK293T cells or *NSUN2^−/−^* HEK293T cells, with or without infection by SeV for 8 h. (f) qPCR analysis of *VSV-G* RNA and VSV plaque assay in wild-type HEK293T cells or *NSUN2^−/−^* HEK293T cells with infection by VSV-GFP for 24 h (MOI = 0.005). (g) Microscopy analysis of VSV-GFP replication in wild-type HEK293T cells or *NSUN2^−/−^* HEK293T cells, both infected with VSV-GFP for 24 h (MOI = 0.005). (h) qPCR analysis of *VSV-G* RNA in wild-type HEK293T cells or *NSUN2^−/−^* HEK293T cells with infection by VSV-GFP for 24 h (MOI = 0.005), with or without ruxolitinib treatment. (i) qPCR analysis of *Nsun2* mRNA in bone-marrow-derived dendritic cells (BMDCs) from 8-week-old wild-type C57BL/6 mice with infection by SeV, HSV-1, VSV, or ZIKV for 0, 24, 48, and 72 h. (j) qPCR analysis of *NSUN2* mRNA in Caco-2 cells with infection by SARS-CoV-2 for 0, 4, 12, and 24 h (MOI = 0.1). (k) RNA-seq signals for *NSUN2* in bronchoalveolar lavage fluid (BALF) of COVID-19 patients (Patient1, Patient2) and healthy controls (Ctrl1, Ctrl2, Ctrl3). Total RNA was extracted and analyzed by RNA-seq to identify differentially expressed genes implicated in COVID-19 disease pathogenesis. The scale on the y-axis indicates the read density per million of total normalized reads. Data are representative of three independent experiments and were analyzed by two-tailed unpaired t test. Graphs show the mean ± SD (n = 3) derived from three independent experiments. NS, not significant for *P* > 0.05, \**P* < 0.05, \*\**P* < 0.01, \*\*\**P* < 0.001.

We next investigated whether NSUN2 is involved in antiviral responses during vesicular stomatitis virus (VSV) infection. Knockout of NSUN2 in HEK293T significantly inhibited the replication of VSV carrying a green fluorescent protein (GFP) reporter (VSV-GFP) (**Fig. 1f and 1g**). The same results were also obtained in NSUN2-knockout A549 cell lines compared with wild-type A549 cells (**Supplementary Fig S1b-e**). These results indicate that knockout of NSUN2 results in cells being less vulnerable to VSV-GFP infection compared to wild-type cells. To further confirm that the inhibition of VSV replication in the NSUN2-deficient cells was indeed due to more potent type I interferon responses, we tested whether inhibition of interferon pathway affected VSV propagation. For this, we used ruxolitinib, a potent and selective JAK 1/2 inhibitor that blocks signaling downstream of type I interferon receptors. As shown in **Fig. 1h**, the inhibition of VSV propagation in NSUN2-knockout cells could be rescued by ruxolitinib treatment, which further confirms that the effects of NSUN2 deficiency on VSV propagation involve antiviral type I interferon responses. These results strongly suggest that NSUN2 is a negative regulator of type I interferon responses and that NSUN2 deficiency prominently enhances antiviral innate responses and, thus, inhibits virus propagation.

To further investigate the biological role of NSUN2 during viral infection, we observed that the *Nsun2* mRNA indeed decreased with the progression of time following infection of bone-marrow-derived dendritic cells (BMDCs) by SeV, herpes simplex virus 1 (HSV-1), VSV, or Zika virus (ZIKV), which reveals the potential role of NSUN2 during viral infections (**Fig. 1i**). Of note, we found that SARS-CoV-2 infection could also significantly reduce *NSUN2* mRNA levels in Caco-2 cells (**Fig. 1j**). We further carried out transcriptome sequencing of the RNAs isolated from the bronchoalveolar lavage fluid (BALF) of two COVID-19 patients (39). *NSUN2* mRNA was consistently reduced in COVID-19 patients compared with healthy individuals (**Fig. 1k**). Taken together, the results indicate that NSUN2 may serve as a negative regulator of type I interferon responses, and that expression of NSUN2 is dramatically reduced to enhance antiviral type I interferon responses during infection with different viruses, including SARS-CoV-2.

### NSUN2 inhibits type I interferon responses by regulating IRF3 expression levels

To investigate the mechanism of NSUN2 in the regulation of type I interferon responses, exogenous NSUN2 expression markedly suppressed the PRDIII-I-luc activity induced by upstream activators, including RIG-I, MDA5, MAVS, TBK1, and the constitutively active phosphorylation mimetic IRF3-5D (**Fig. 2a**), while knockdown of NSUN2 had the opposite effect (**Fig. 2b**). Since IRF3 is the final factor in the process of initiation of type I interferon responses, we speculated that NSUN2 may exert its function at IRF3 node. As shown in **Fig. 2c**, immunoblot analysis revealed that exogenous NSUN2 expression could inhibit the expression of endogenous IRF3 and that the levels of endogenous IRF3 were enhanced in NSUN2-knockout cells compared with those in wild-type HEK293T (**Fig. 2d**) and A549 (**Fig. 2e**) cells. By contrast, endogenous TBK1 protein levels did not show significant change. Moreover, knockout of NSUN2 promoted levels of IRF3 Ser396 phosphorylation but not TBK1 Ser172 phosphorylation (**Fig. 2e and 2f**). These results demonstrate that NSUN2 deletion could enhance the overall levels of IRF3 protein as well as its phosphorylation. Next, we conjugated the IRF3-CDS (coding sequence) with EGFP to allow for visual characterization of responses in IRF3 expression by fluorescence. We observed that exogenous NSUN2 expression inhibited the fluorescence of IRF3-CDS-EGFP (**Fig. 2g**). To summarize, these results reveal that NSUN2 could specifically inhibit the expression of IRF3 and thus negatively regulate type I interferon responses following virus infection.

**Figure 2.**
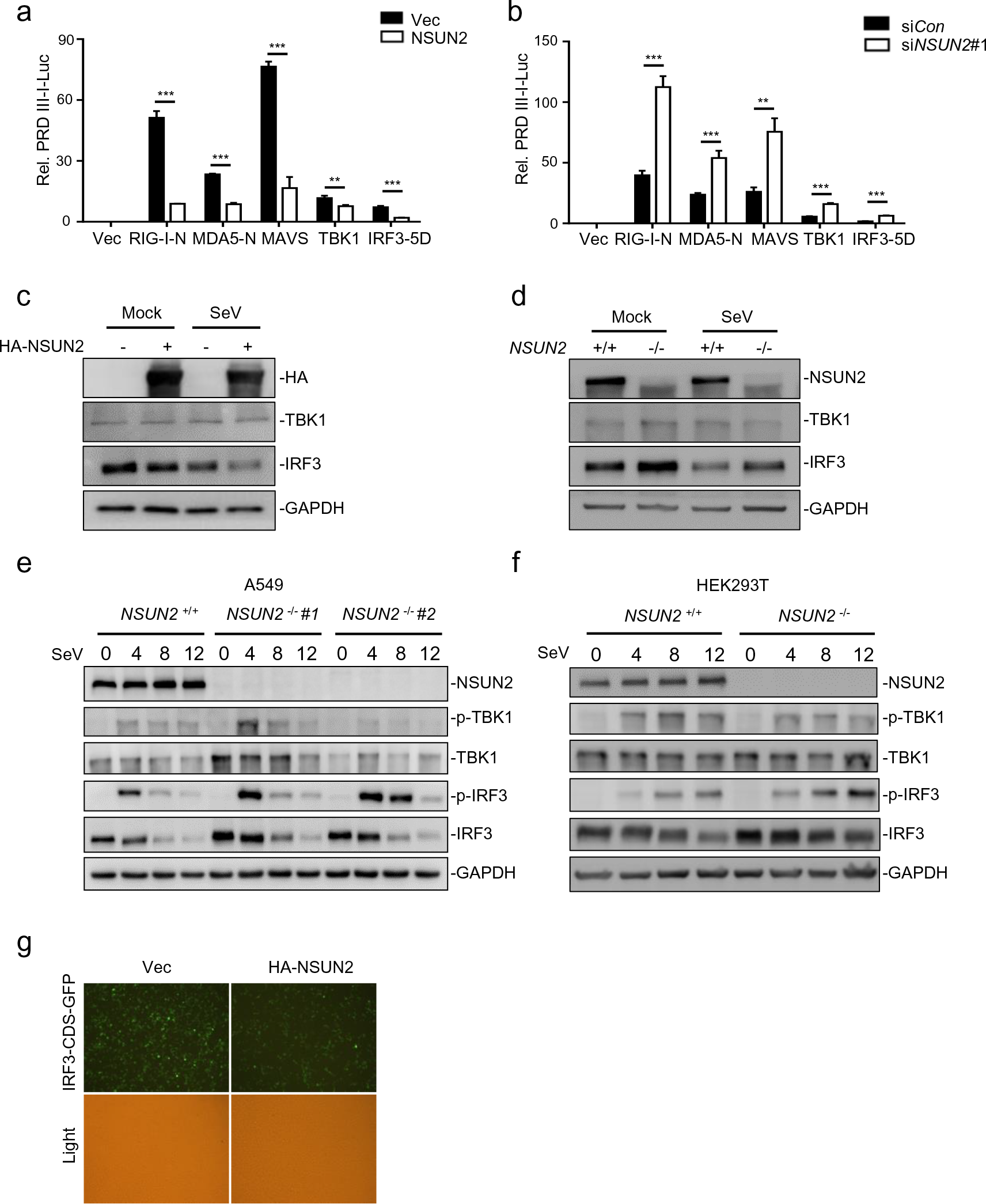
NSUN2 inhibits the expression level of IRF3. (a) Dual-luciferase assay analyzing a luciferase reporter plasmid for the IRF3-responsive promoter containing positive regulatory domains III and I of the IFN-β promoter (PRDIII-I-Luc) in HEK293T cells in 24-well plates transfected for 36 h with the RIG-N, MDA5-N, MAVS, TBK1, and IRF3-5D expression plasmids, as indicated, with co-transfection with empty vector or NSUN2. (b) Dual-luciferase analysis of PRDIII-I-Luc in HEK293T cells in 24-well plates transfected for 36 h with the indicated RIG-N, MDA5-N, MAVS, TBK1, and IRF3-5D expression plasmids with co-transfection with siControl or siNSUN2-1. (c) Immunoblot analysis in HEK293T cells transfected with vector or NSUN2 for 36 h, with or without infection by SeV for another 12 h. (d) Immunoblot analysis in wild-type HEK293T cells or *NSUN2^−/−^* HEK293T cells with or without infection by SeV for 12 h. (e) Immunoblot analysis in wild-type A549 cells or *NSUN2^−/−^* A549 cells with infection by SeV for 0, 4, 8, and 12 h. (f) Immunoblot analysis in wild-type HEK293T cells or NSUN2*^−/−^* HEK293T cells, with infection by SeV for 0, 4, 8, and 12 h. (g) Immunofluorescence microscopy of HEK293T cells transfected with IRF3-CDS-EGFP along with vector or NSUN2 for 36 h. Data are representative of three independent experiments and were analyzed by two-tailed unpaired t test. Graphs show the mean ± SD (n = 3) derived from three independent experiments. NS, not significant for *P* > 0.05, \**P* < 0.05, \*\**P* < 0.01, \*\*\**P*-< 0.001.

### NSUN2 catalyzes m^5^C methylation of *IRF3* mRNA

Since NSUN2 has been reported to regulate some genes by methylating their mRNAs and affecting RNA fate or function (17, 19, 20), we speculated that it might physically interact with *IRF3* mRNA. Firstly, co-immunoprecipitation followed by immunoblot analysis showed that there was no interaction between NSUN2 and IRF3 protein in HEK293T (**Fig. 3a**). We further overexpressed and immunoprecipitated NSUN2 protein in SeV-stimulated HEK293T cells and subjected it to RNA extraction and qPCR. The results reveal that NSUN2 indeed binds with endogenous *IRF3* mRNA, while endogenous *TBK1* mRNA did not interact with NSUN2 (**Fig. 3b**). Furthermore, knockdown or knockout of NSUN2 boosted endogenous *IRF3* mRNA levels while endogenous *TBK1* mRNA levels were not affected (**Fig. 3c and 3d, upplementary Fig S2**). We then detected the half-life of endogenous *IRF3* mRNA in wild-type and *NSUN2^−/−^* HEK293T cells following treatment of actinomycin D (ActD) which inhibits mRNA transcription in mammalian cells. The results show that knockout of NSUN2 significantly increased the half-life of *IRF3* mRNA from 6.48 h in wild-type cells to 12.39 h in NSUN2 knockout cells (**Fig. 3e**), while the half-life of *TBK1* mRNA had no significant difference, from 4.48 h in wild-type cells to 5.08 h in NSUN2 knockout cells. Consistent results were also found in A549 cells, as shown in **Supplementary Fig S3**. These results indicate that NSUN2 decreased IRF3 protein levels dramatically by binding to *IRF3* mRNA and accelerating its degradation.

**Figure 3.**
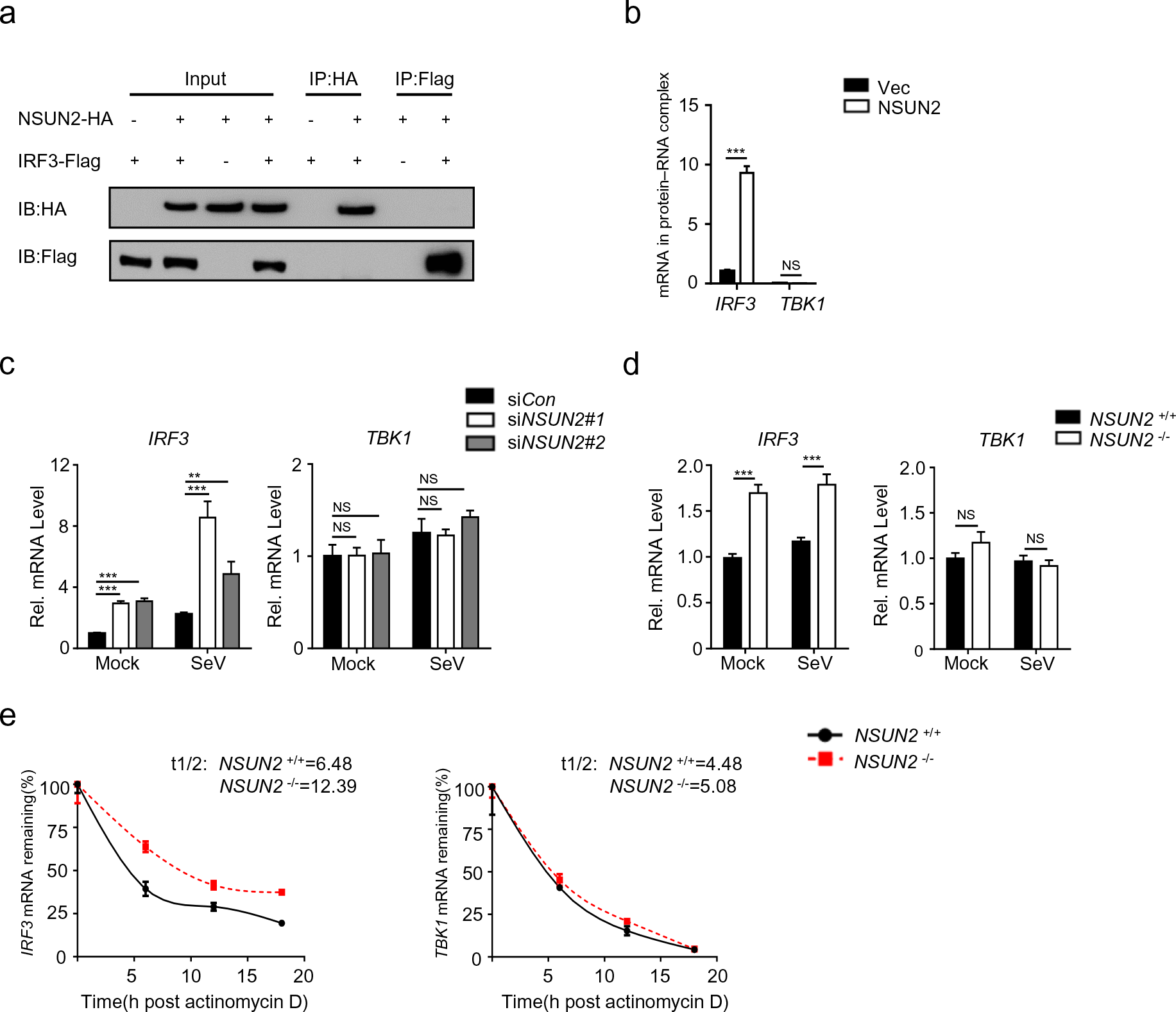
NSUN2 interacts with *IRF3* mRNA and induces its degradation. (a) Coimmunoprecipitation (IP) and immunoblot (IB) analysis of HEK293T cells transfected with plasmids encoding HA-NSUN2 and Flag-IRF3. (b) Immunoprecipitation by HA-Tag-conjugated beads and immunoblot analysis of HEK293T cells transfected with plasmids encoding HA-NSUN2, with SeV infection for 8 h, followed by RNA extraction and qPCR analysis of combined *IRF3* mRNA. (c) qPCR analysis of *IRF3* mRNA and *TBK1* mRNA in HEK293T cells transfected with siControl or siRNAs targeting NSUN2, with or without infection by SeV, for 8 h. (d) qPCR analysis of *IRF3* mRNA and *TBK1* mRNA in wild-type HEK293T cells or *NSUN2^−/−^* HEK293T cells, with or without infection by SeV for 8 h. (e) Stability analysis of *IRF3* mRNA and *TBK1* mRNA in wild-type HEK293T cells or *NSUN2^−/−^* HEK293T cells with treatment of actinomycin D (ActD) for 0, 6, 12, and 18 h. Data are representative of three independent experiments and were analyzed by two-tailed unpaired t test. Graphs show the mean ± SD (n = 3) derived from three independent experiments. NS, not significant for *P* > 0.05, \**P* < 0.05, \*\**P* < 0.01, \*\*\**P* < 0.001.

Since NSUN2 is a typical RNA methyltransferase catalyzing the formation of m^5^C in coding and non-coding RNAs, we speculated that NSUN2 might catalyze the formation of m^5^C in *IRF3* mRNA and then induce its degradation. Therefore, we prepared RNA segments of *RIG-I*, *MAVS*, *TBK1*, and *IRF3*, the four key signaling molecules that determine the innate immune response to viral infection, by *in vitro* transcription. Micro-125b, which can be methylated by NSUN2, served as a positive control (40). The RNAs were used for *in vitro* methylation assays using recombinant GST-NSUN2 and ^3^H-labeled S-adenosyl methionine (SAM). The transcribed *IRF3* mRNA could be highly methylated by NSUN2 compared with transcripts of *RIG-I*, *MAVS*, and *TBK1* **(Fig. 4a and 4b)**. The data suggest that NSUN2 could specifically mediate the methylation of *IRF3* mRNA *in vitro*. To determine which region might be methylated, we divided *IRF3* mRNA into seven parts, including 5’UTR (1–235 nt), CDS1 (236–485 nt), CDS2 (486–735 nt), CDS3 (736–985 nt), CDS4 (986–1235 nt), CDS5 (1236–1519 nt), and 3’UTR (1520–1595 nt) (**Fig. 4a**). As is demonstrated in **Fig. 4c**, *IRF3* 5’UTR, 3’UTR, CDS2, and CDS3 were highly methylated by NSUN2 compared with other segments. To further verify whether endogenous *IRF3* mRNA could be methylated by NSUN2 *in vivo*, we pulled down endogenous *IRF3* mRNA using specific *IRF3* CHIRP probes which were 3’biotin-TEG-modified. Equal amounts of endogenous *IRF3* mRNA were loaded on the membrane, and the levels of m^5^C were assayed. As is shown in **Fig. 4d**, the m^5^C methylation level of *IRF3* mRNA in NSUN2 knockout cells was markedly lower than that of wild-type cells. Reconstitution of exogenous NSUN2 into NSUN2 knockout cells restored the m^5^C methylation levels of endogenous *IRF3* mRNA. Consistent with this, the results of m^5^C MeRIP showed that the levels of endogenous m^5^C methylated *IRF3* mRNA in NSUN2 knockout cells was significantly lower than in wild-type cells, and exogenous NSUN2 expression could dramatically enhance the levels of endogenous m^5^C-methylated *IRF3* mRNA (**Fig. 4e**). To investigate the biological function of m^5^C methylation of *IRF3* mRNA by NSUN2, we constructed pGL3-derived reporters bearing either IRF3-5’UTR, IRF3-CDS, or IRF3-3’UTR. We tested the activity of these reporters in NSUN2-knockout HEK293T cells compared with those in wild-type HEK293T. As shown in **Fig. 4f**, knockout of NSUN2 could significantly increase the luciferase activity of reporter pGL3-IRF3-5’UTR, pGL3-IRF3-CDS, and pGL3-IRF3-3’UTR. The above results demonstrate that NSUN2 could mediate m^5^C methylation of *IRF3* mRNA both *in vitro* and *in vivo*, and that the four highly methylated regions in *IRF3* mRNA are the major targets of NSUN2. This methylation might result in the degradation of *IRF3* mRNA and, thereby, decreased levels of IRF3 protein.

**Figure 4.**
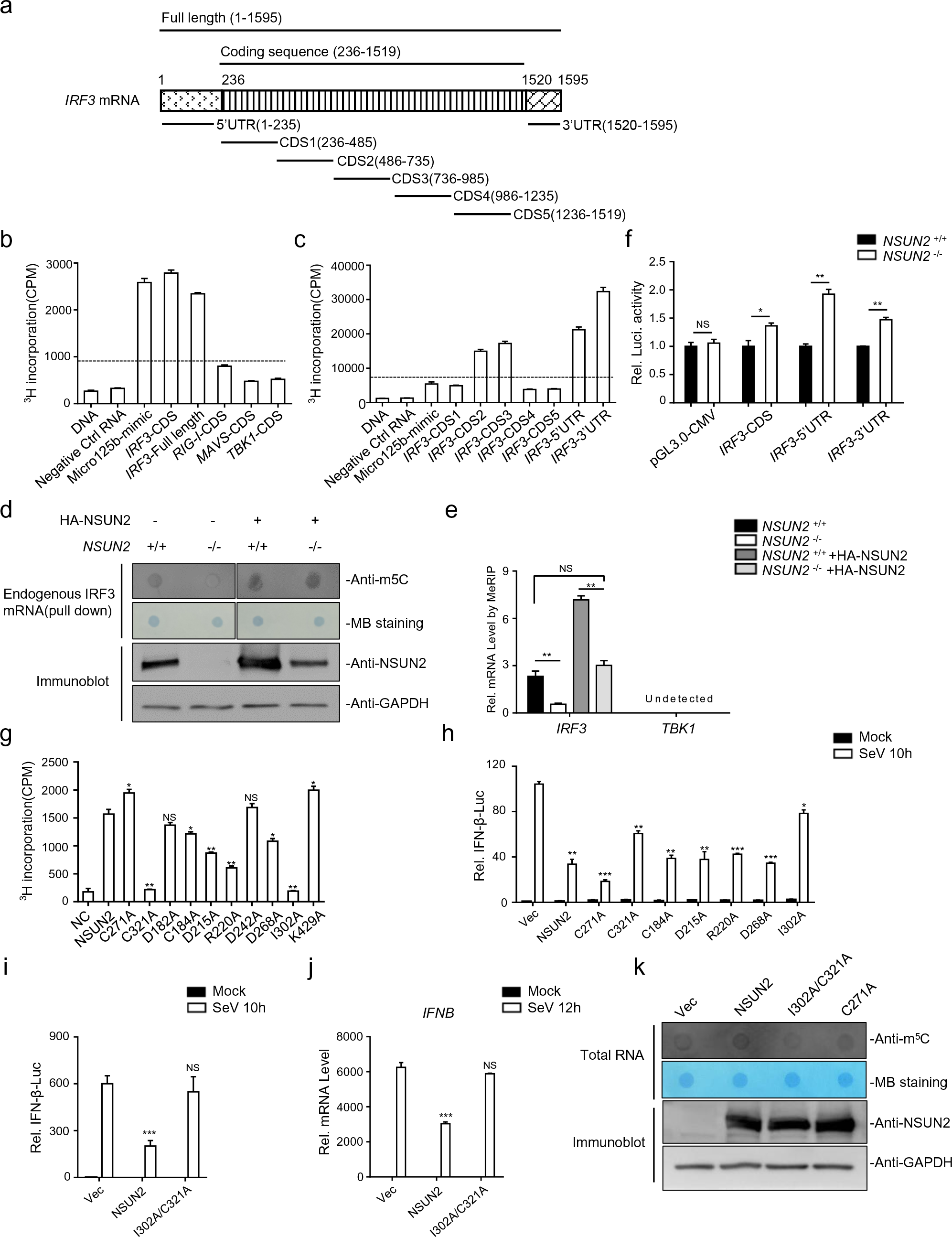
NSUN2 catalyzes the formation of m^5^C methylation of *IRF3* mRNA both exogenously and endogenously. (a) Schematic diagram of the *IRF3* mRNA segments used for *in vitro* methylation assays and bisulfite RNA sequencing. (b) *In vitro* m^5^C methylation assays using recombinant GST-NSUN2 and the *in vitro* transcripts. (c) *In vitro* m^5^C methylation assays using recombinant GST-NSUN2 and the *in vitro* transcribed segments of *IRF3* mRNA depicted in Figure 4a. (d) m^5^C dot blot analysis of endogenous *IRF3* mRNA (200 ng) pulled down by IRF3 CHIRP probes in wild-type HEK293T cells or *NSUN2^−/−^* HEK293T cells with or without exogenous NSUN2 overexpression. Equal *IRF3* mRNAs were also loaded and verified by methylene blue (MB) staining. (e) MeRIP analysis of the m^5^C methylated *IRF3* mRNA immunoprecipitated by m^5^C antibody from wild-type HEK293T cells or *NSUN2^−/−^* HEK293T cells, with or without exogenous NSUN2 expression. *TBK1* was used as a negative control. (f) Wild-type HEK293T cells or *NSUN2^−/−^* HEK293T cells were transfected with pGL.3.0-CMV-Luc or pGL3.0-CMV-*IRF3*-CDS-Luc or pGL3.0-CMV-*IRF3*-5’UTR-Luc or pGL3.0-CMV-*IRF3*-3’UTR-Luc, together with *Renilla* luciferase (RL-TK). Forty-eight hours later, firefly luciferase activity against *Renilla* luciferase activity was analyzed. (g) In vitro m^5^C methylation assays using recombinant GST-NSUN2 and different mutant proteins. (h-i) Dual-luciferase assay analyzing IFN-β promoter activity in HEK293T cells (h) or *NSUN2^−/−^* HEK293T cells (i) in 24-well plates transfected for 24 h with 100 ng IFN-β firefly luciferase reporter (IFN-β-Luc) and 20 ng *Renilla* luciferase (RL-TK), along with 300 ng vector or plasmid encoding NSUN2 or different mutants, with or without infection by SeV, for another 10 h. (j) qPCR analysis of *IFNB* mRNA in *NSUN2^−/−^* HEK293T cells transfected for 24 h with NSUN2 or different mutants, with or without infection by SeV, for another 12 h. (k) m^5^C dot blot analysis of total RNA (1 µg) extracted from in *NSUN2^−/−^* HEK293T cells with exogenous NSUN2 expression or different mutants. Equal RNAs were also loaded and verified by methylene blue (MB) staining. Data are representative of three independent experiments and were analyzed by two-tailed unpaired t test. Graphs show the mean ± SD (n = 3) derived from three independent experiments. NS, not significant for *P* > 0.05, \**P* < 0.05, \*\**P* < 0.01, \*\*\**P* < 0.001.

To further confirm whether m^5^C methyltransferase activity of NSUN2 is the determining factor that results in the inhibition of interferon responses, we generated different NSUN2 methyltransferase mutants, including C271A and C321A, which are reported to be the key sites whereby their mutation may inhibit NSUN2 m^5^C methyltransferase activity (22), as well as several predicted inactiving mutations. The *in vitro* methylation results show that the NSUN2 mutants, including C184A, D215A, R220A, and D268A, had partially decreased methylation activity, while C321A and I302A mutations almost completely abolished catalytic activity. However, C271A resulted in mildly increased catalytic activity of NSUN2 (**Fig. 4g**). Of note, we investigated the relationship between the methylation activities and the stimulation of IFN-β pathway in an SeV-triggered IFN-β-Luc reporter system. As shown in **Fig. 4h**, some of inhibition ability in SeV-induced-IFN-β luciferase assay was lost following overexpression of either I302A or C321A compared with wild-type NSUN2, while C271A could enhance the inhibition ability compared with wild-type NSUN2. Moreover, we found that the double mutant I302A/C321A had totally lost its inhibition ability in terms of both function (**Fig. 4i**) and effects on *IFNB* mRNA levels (**Fig. 4j**) in *NSUN2^−/−^* HEK293T cells. We further detected the m^5^C methylation levels of total RNA in *NSUN2^−/−^* HEK293T cells transfected with NSUN2 or its mutants using dot blot analysis. In accordance with the above results, I302A/C321A double mutation resulted in almost complete loss of m^5^C methyltransferase activity, while C271A still maintained m^5^C methyltransferase activity (**Fig. 4k**). Moreover, ALYREF has been characterized as an m^5^C reader in the nucleus, facilitating the export of m^5^C-modified mRNAs (22). The negatively regulation of type I interferon responses by NSUN2 may also occur in collaboration with ALYREF, which has also been independently observed to negatively regulate type I interferon responses, as depicted in **Supplementary Fig S4**. To summarize, NSUN2 could catalyze the formation of m^5^C modification of *IRF3* mRNA and accelerate its fast turnover and regulate IRF3-mediated type I interferon responses. Of note, this regulation by NSUN2 is dependent on its m^5^C methyltransferase activity.

### Four methylated cytosines of *IRF3* mRNA were identified to regulate RNA levels

We also aimed to identify the exact methylation cytosines in *IRF3* mRNA. Using bisulfite sequencing assays (**Fig. 5a**), we identified four cytosines in *IRF3* mRNA as major sites of methylation that were highly methylated by recombinant NSUN2 protein *in vitro*: C169 (11/20, 55%) in 5’UTR, C1569 (15/20, 75%) in 3’UTR, C556 (9/16, 56.25%) in CDS2 (486–735), and C815 (7/16, 43.75%) in CDS3 (736–985), which is consistent with the four high methylation regions observed earlier in *IRF3* mRNA **(Fig. 4c and 5b-c, Supplementary Fig S5)**. We then tested whether these four identified highly methylated cytosines are indeed methylated and involved in the regulation of *IRF3* mRNA by NSUN2 protein. It was observed that mutations C169nt (C to G) in the 5’UTR, C1569nt (C to G) in the 3’UTR, C556nt (C to T) in CDS2 (486–735) and C815nt (C to A) in CDS3 (736–985) reduced the methylation level by half in biochemical assays with recombinant NSUN2 (**Fig. 5d**). We then constructed expression plasmids containing either wild type IRF3 full length (IRF3-FL, 1-1595nt) or various site-mutated IRF3-FLs. We observed that mutations of the four cytosines could consistently enhance the expression levels of *IRF3* mRNA in *Irf3^−/−^Irf7^−/−^* MEFs compared to wild type IRF3 full length (IRF3-FL-WT) (**Fig. 5e**). Correspondingly, the IRF3-mediated *Ifnb* mRNA levels were also remarkably elevated upon SeV infection (**Fig. 5f**). We also utilized the lentiviral system to generate stable IRF3 cell lines in *Irf3^−/−^Irf7^−/−^* MEFs. The *IRF3* mRNA levels in the IRF3-FL-Mut (IRF3-FL-5’&3’UTR-CDS2&3-Cm) stable cell line was 15-fold higher than that in the IRF3-FL-WT stable cell line. Moreover, exogenous NSUN2 expression significantly reduced *IRF3* mRNA levels in the IRF3-FL-WT stable cell line, while *IRF3* mRNA levels in the IRF3-FL-Mut stable cell line were mildly decreased, indicating that methylation of these four cytosines in *IRF3* mRNA might predominantly influence its stability (**Fig. 5g**). To confirm this, we measured the stability of these transcripts and found that the IRF3-FL-Mut transcript was remarkably more stable than the IRF3-FL-WT transcript, indicating that methylation of these four cytosines by NSUN2 is indeed critical for regulating *IRF3* mRNA stability (**Fig. 5h**). Taken together, our results demonstrate that the loss of m^5^C modification could lead to increased stability of *IRF3* mRNA and enhanced IFN-β production, thus facilitating a stronger antiviral response, and that the four highly methylated cytosines in *IRF3* mRNA play a critical role in NSUN2-mediated regulation of antiviral responses.

**Figure 5.**
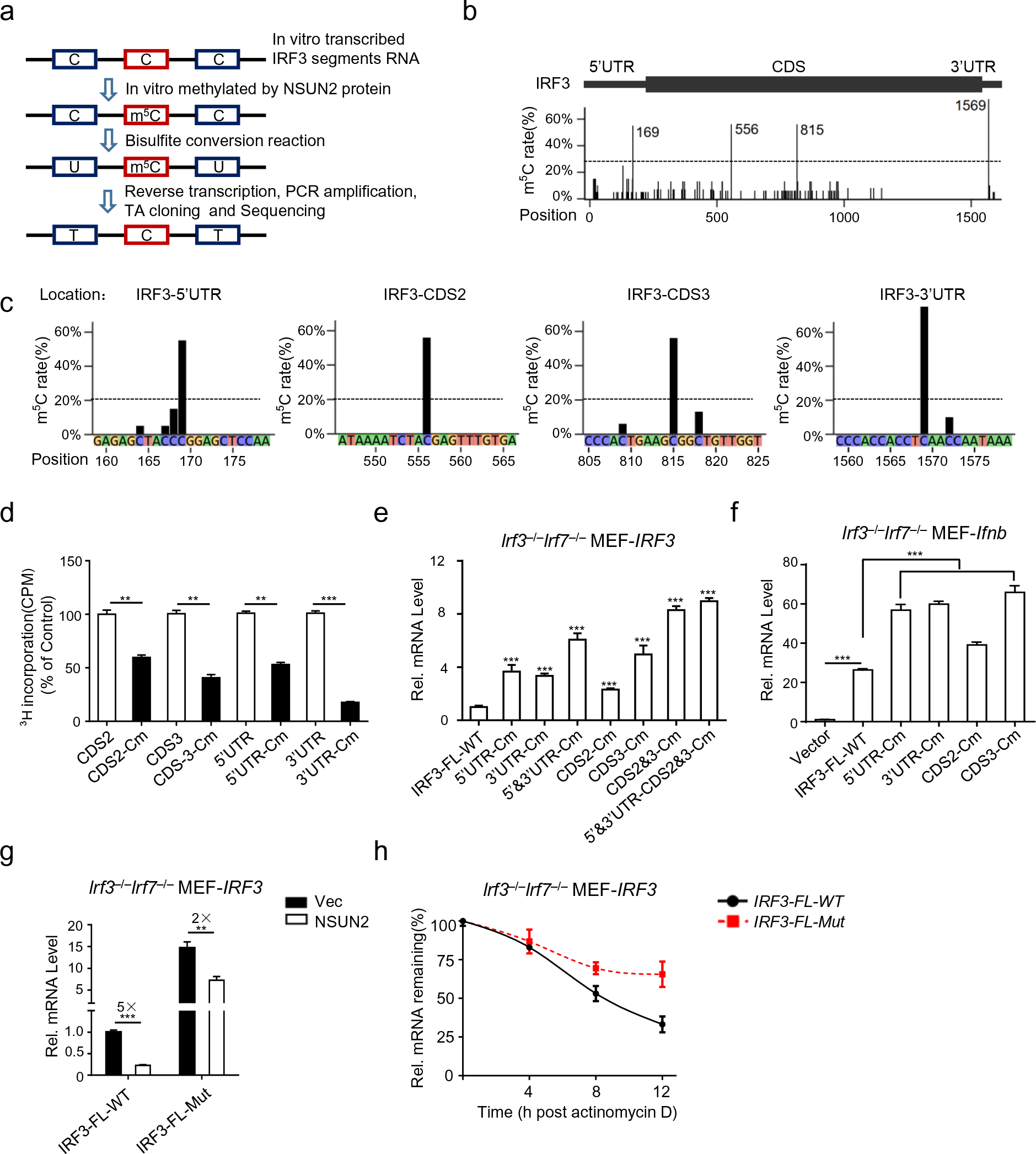
*IRF3* m^5^C methylation site mutation results in enhanced *IRF3* expression and antiviral response. (a) Schematic depiction of *in vitro* bisulfite RNA sequencing to distinguish m^5^C (cytosine methylated by NSUN2) from C (cytosine not methylated). (b-c) Identification of m^5^C modification on all cytosines of *IRF3* mRNA. Data are expressed as the ratio of m^5^C to (C + m^5^C). (d) *In vitro* m^5^C methylation assays of the *IRF3* segments or the m^5^C methylated cytosines mutated segments using recombinant GST-NSUN2. (e) qPCR analysis of *IRF3* mRNA in *Irf3*^−/−^*Irf7*^−/−^ MEFs transfected with plasmid encoding NSUN2 along with wild-type IRF3 full length (IRF3-FL-WT) or various cytosine-mutated IRF3-FLs for 48 h. (f) qPCR analysis of *Ifnb* mRNA in *Irf3*^−/−^*Irf7*^−/−^ MEFs transfected with plasmid encoding NSUN2 along with IRF3-FL-WT or the m^5^C methylated cytosines mutated IRF3-FL, with stimulation by SeV, for 8 h. (g) qPCR analysis of *IRF3* mRNA in *Irf3*^−/−^*Irf7*^−/−^ MEFs reconstituted with IRF3-FL-WT or IRF3-FL-Mut (IRF3-FL with the four m^5^C methylated cytosines mutated) by lentiviral system transfected with plasmid encoding NSUN2 or empty vector (Vec). (h) Stability analysis of *IRF3* mRNA in *Irf3*^−/−^*Irf7*^−/−^ MEFs reconstituted with IRF3-FL-WT or IRF3-FL-Mut by lentiviral system with treatment of actinomycin D (ActD) for 0, 4, 8, and 12 h. Data are representative of three independent experiments and analyzed by two-tailed unpaired t test. Graphs show the mean ± SD (n = 3) derived from three independent experiments. NS, not significant for *P* > 0.05, \**P* < 0.05, \*\**P* < 0.01, \*\*\**P* < 0.001.

### Pivotal role of NSUN2 in the induction of type I interferon and antiviral response *in vivo*

To determine the role of NSUN2 in antiviral response *in vivo*, we created targeted deletions of NSUN2 in mice by removing 10 bp in exon 3 of *Nsun2* genome by CRISPR–Cas9, which resulted in a frameshift mutation. However, we found that *Nsun2^−/−^* mice died in utero. We found that *Nsun2^+/−^* progeny could reach adulthood, so we chose *Nsun2^+/−^* mice as “NSUN2-knockdown mice”. As expected, the *Nsun2* expression in *Nsun2^+/−^* mice did reduce by half than their wild-type littermates (**Fig. 6a**). We then investigated innate antiviral responses in *Nsun2^+/+^* mice and *Nsun2^+/−^* mice. As shown in **Fig. 6b**, the production of *Ifnb* mRNA was more dramatically enhanced in bone-marrow-derived dendritic cells (BMDCs) from *Nsun2^+/−^* mice than in those from their wild-type littermates following infection with SeV, HSV-1 or VSV. The IFN-β mediated downstream *Isg15* and *Cxcl10* were also significantly enhanced in BMDCs from *Nsun2^+/−^* mice (**Fig. 6c**). We also observed significantly higher IFN-β and IFN-α production in the serum of *Nsun2^+/−^* mice after intraperitoneal injection of VSV by ELISA (**Fig. 6d**). Furthermore, we found a higher IFN-β production and a lower viral burden of VSV in various organs of *Nsun2^+/−^* mice than in wild-type mice at the mRNA levels (**Fig. 6e**). We also compared the survival rates after intraperitoneal injection of VSV. The results indicate that *Nsun2^+/+^* mice were more vulnerable to VSV-triggered mortality than were *Nsun2^+/−^* mice (**Fig. 6f**). All these data suggest that NSUN2, the expression of which is reduced during viral infection, was quite pivotal for the induction of type I interferon and antiviral responses *in vivo*.

**Figure 6.**
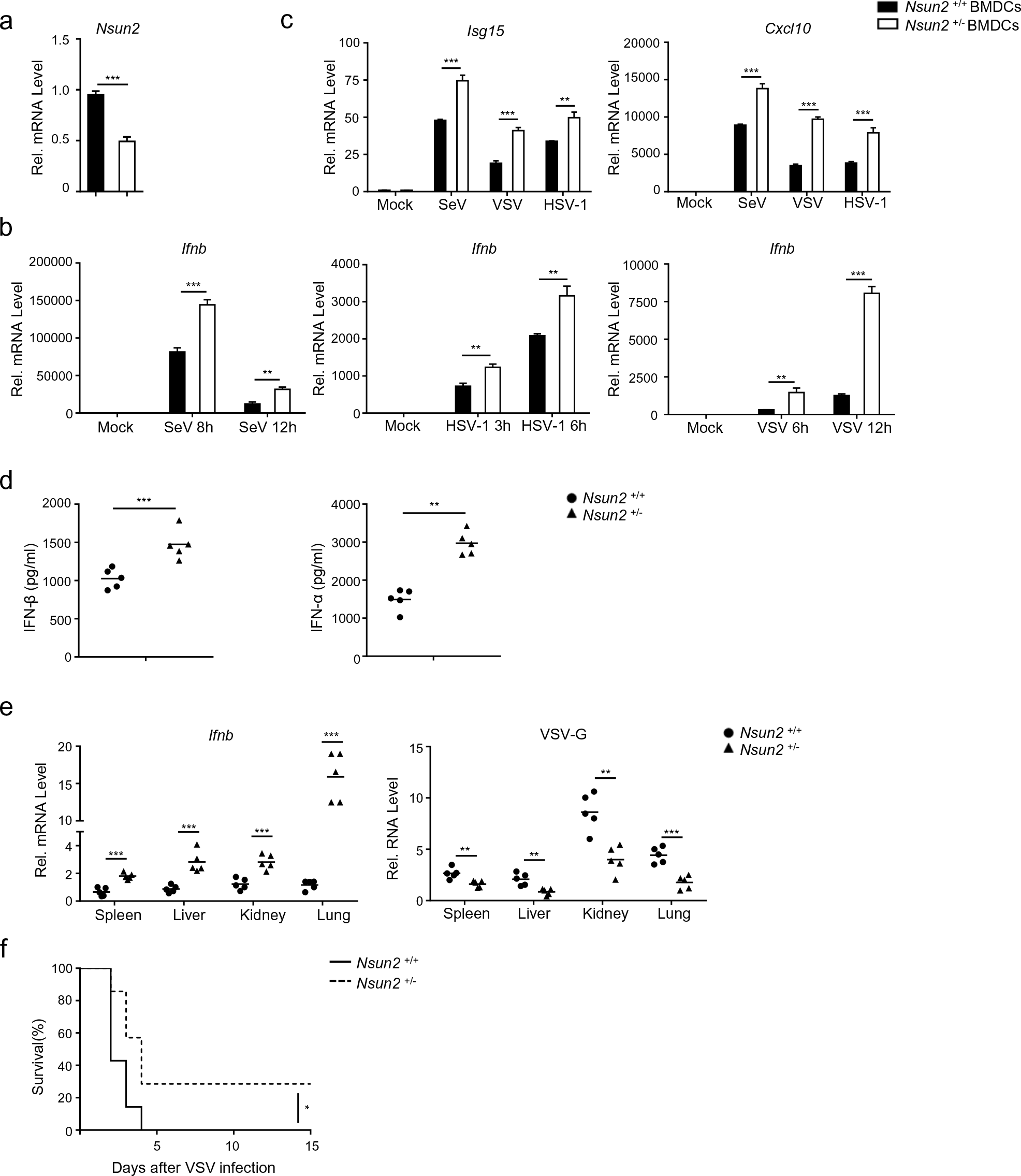
Pivotal role for NSUN2 in the induction of type I interferon and antiviral response *in vivo*. (a) qPCR analysis of *Nsun2* mRNA in BMDCs from *Nsun2^+/+^* mice or *Nsun2^+/−^* mice. (b) qPCR analysis of *Ifnb* mRNA in BMDCs from *Nsun2^+/+^* mice or *Nsun2^+/−^* mice, with or without infection by SeV for 8 and 12 h, HSV-1 for 3 and 6 h, or VSV for 6 and 12 h. (c) qPCR analysis of *Isg15* and *Cxcl10* in BMDCs from *Nsun2^+/+^* mice or *Nsun2^+/−^* mice, with or without infection by SeV, HSV-1, or VSV. (d) ELISA of IFN-β and IFN-α in serum from 8-week-old *Nsun2^+/+^* mice (n = 5) and *Nsun2^+/−^* mice (n = 5) injected intraperitoneally for 16 h with VSV (4 × 10^7^ PFU per mouse). Each symbol represents an individual mouse; small horizontal lines indicate the mean. (e) qPCR analysis of *Ifnb* mRNA and the corresponding *VSV-G* RNA in different organs from *Nsun2^+/+^* mice or *Nsun2^+/−^* mice, injected intraperitoneally for 16 h with VSV (4 × 10^7^ PFU per mouse). (f) Survival (Kaplan–Meier curve) of *Nsun2^+/+^* mice (n = 7) or *Nsun2^+/−^* mice (n = 7) infected intraperitoneally with a high dose of VSV (1 × 10^8^ PFU per mouse) and monitored for survival for 15 days. Data are representative of three independent experiments and were analyzed by two-tailed unpaired t test. Graphs show the mean ± SD (n = 3) derived from three independent experiments. NS, not significant for *P* > 0.05, \**P* < 0.05, \*\**P* < 0.01, \*\*\**P* < 0.001.

## Discussion

Antiviral innate immunity involves sophisticated signaling pathways for sensing pathogens and initiating innate immune responses against infection, which requires ingenious regulation at different levels including transcriptional, translational, and post-translational. It is known that IRF3, which plays a vital role in the initiation of type I interferon responses after infection, is regulated by multiple modifications, such as phosphorylation, ubiquitination, and acetylation, which function in maintaining immune homeostasis (38, 41, 42). Recently, the m^6^A machinery has been reported to be involved in immune responses via epigenetic modification. For example, it has been reported that the m^6^A machinery could inhibit the innate immune response to infection by directly dictating the fast turnover of *IFNB* mRNAs and consequently facilitating viral propagation (10). Another study demonstrated that ALKBH5 could erase the m^6^A modification of *MAVS*, *TRAF3*, and *TRAF6* mRNAs, enforce their retention in the nucleus and result in their decreased translation and inhibited type I interferon production (13). Moreover, hnRNPA2B1 was reported to function as an m^6^A “modulator” that promotes m^6^A modification and nucleocytoplasmic trafficking of *CGAS*, *IFI16*, and *STING* mRNAs in response to DNA virus infection, leading to the enhanced production of type I interferons (14). The effects of m^6^A modification on interferon responses may vary because of the different systems and different readers and precise downstream regulation. However, no report has demonstrated that m^6^A modification could regulate interferon responses by directly methylating IRF3. More importantly, there are, to date, no reports of m^5^C modification regulating antiviral innate immunity.

In this study, we revealed a novel mechanism by which the m^5^C machinery functions in innate immune responses via the methylation of *IRF3* mRNA to negatively regulate type I interferon responses, indicating that the m^5^C and m^6^A machineries may have different specificities with respect to regulating multiple signaling molecules involved in antiviral innate immune responses. We demonstrated that NSUN2 could specifically methylate *IRF3* mRNA via four major cytosine sites. The mutation of these four major cytosines enhanced the stability and expression of *IRF3* mRNA (**Fig. 5**) and, thereby, interferon responses. Moreover, in our system, the m^6^A machinery was also found to be involved in regulating interferon responses (**Fig. 1a**), but the overall effect was not significant compared with the m^5^C machinery, which may be because the m^6^A machinery regulates other signaling molecules with different effects, as mentioned above. However, we do not preclude the possibility that other mechanisms beyond an elevation in *IRF3* mRNA stability may contribute to the stronger type I interferon responses following knockout of NSUN2. We may speculate that the mRNAs of some other signaling molecules, or the *IFNB* mRNA, may also be m^5^C-modified by NSUN2, such as is the case with m^6^A modification. Future work is required to demonstrate how m^5^C methylation and its downstream recognition and regulation collaboratively and precisely function in antiviral innate immunity.

Moreover, we found that the regulation of type I interferon responses by NSUN2 was dependent on its m^5^C methyltransferase activity. According to our results (**Fig. 4g–k**), the NSUN2 I302A/C321A mutant had almost completely lost its m^5^C methyltransferase activity and ability to regulate type I interferon responses, which is in contrast with the reports of C271A/C321A mutation of NSUN2 (22, 43). In our study, the C271A mutation maintained m^5^C methyltransferase activity in biochemical assays and could still negatively regulate interferon responses. The discrepancy in the key sites of NSUN2 methyltransferase activity may be due to the different systems and the different roles NSUN2 plays in multiple physiological processes. Further work is required to uncover the structure of NSUN2 protein and the key sites that determine its m^5^C methyltransferase activity and regulation activity in multiple physiological processes.

NSUN2 and TRDMT1 (DNMT2) are two m^5^C methyltransferases reported in animals, but the identity of the m^5^C demethylase remains unknown (15, 16). In our study, TRDMT1 did not show significant regulation of interferon responses unlike NSUN2. ALYREF has earlier been characterized as an m^5^C reader in the nucleus involved in facilitating the export of m^5^C-modified mRNAs (22). In our results, exogenous NSUN2 expression could dramatically inhibit IFN-β production, and exogenous ALYREF expression could also (**Fig. 1a-c**, **Supplementary Fig S4)**, which further confirmed that m^5^C modification is indeed involved in regulating type I interferon responses. YBX1 was identified as another m^5^C reader that could maintain the stability of its target mRNA by recruiting ELAVL1 (23). In our study, NSUN2 could directly methylate *IRF3* mRNA and accelerates its degradation, which seems to contradict the function of the NSUN2–YBX1–ELAVL1 axis. These two seemingly opposing mechanisms may uncover the different roles that m^5^C modification play in various biological processes. Different m^5^C readers might have different functions and play different roles. For example, YTH family members have been reported to serve as m^6^A readers that recognize m^6^A-modified RNA and further regulate mRNA splicing, translation, or degradation (44-47). The specific degradation mechanism induced by m^5^C and m^6^A modification has not yet been clarified clearly and requires more investigation. Further work is required to delineate these different mechanisms and the different roles that m^5^C readers play. The m^5^C demethylase, which may maintain balance in the m^5^C modification level in various biological processes, must also be identified.

Furthermore, we found that NSUN2 expression is decreased after infections with different viruses, including SeV (negative-strand RNA virus), HSV-1 (DNA virus), VSV (negative-strand RNA virus), ZIKV (positive-strand RNA virus), and especially SARS-CoV-2 (positive-strand RNA virus, beta-coronavirus). Notably, transcriptome sequencing of the RNAs isolated from the bronchoalveolar lavage fluid (BALF) of two COVID-19 patients revealed that NSUN2 expression was dramatically decreased in COVID-19 patients compared with healthy individuals (**Fig. 1i-k**). We can therefore propose a model whereby NSUN2 is constitutively expressed in resting cells and that IRF3 expression is maintained at a relatively low level. During viral infection, endogenous NSUN2 expression levels decrease via unknown mechanism, which require further investigation for their elucidation, and the IRF3 expression level would therefore be elevated to allow a stronger interferon response and the effective elimination of viruses.

In conclusion, our investigation has revealed a novel and profound role for m^5^C modification in regulating type I interferon responses. We have proposed a crosstalk between m^5^C methylation and antiviral innate immunity, and this might benefit the development of efficient therapeutic interventions for infectious diseases. To move forward, further work is urgently needed to precisely demonstrate how m^5^C methylation is involved in antiviral innate immunity and other physiological processes.

## Materials and Methods

### Viruses, cells, and reagents

SARS-CoV-2 WIV04 (IVCAS 6.7512) was kindly provided by Dr. Zheng-Li Shi. Sendai virus (SeV), herpes simplex virus 1 (HSV-1), and vesicular stomatitis virus carrying a GFP reporter gene (VSV-GFP) were kindly provided by Dr. Hong-Bing Shu. Zika virus (ZIKV) was kindly provided by Dr. Bo Zhang. Vesicular stomatitis virus (VSV) was kindly provided by Dr. Ming-Zhou Chen. Human colorectal adenocarcinoma (Caco-2), HEK293T, HeLa, Vero, and A549 cells were maintained in Dulbecco’s modified Eagle’s medium (DMEM) with 10% fetal bovine serum, 100 U/mL penicillin and 100 µg/mL streptomycin, at 37 °C in 5% CO_2_ incubator. Plasmids were transfected using Lipofectamine 2000 (Invitrogen, 11668027) or Neofect (Neofect, TF201201) following the manufacturer’s instructions, and siRNAs (synthesized by RiboBio company) were transfected using Pepmute (SignaGen, SL100566) following the manufacturer’s instructions. Ruxolitinib and actinomycin D were from MCE (MedChemExpress).

### Mice

*Nsun2^+/−^* C57BL/6J mice were obtained from GemPharmatech Company (Nanjing, China) and housed and bred in specific pathogen-free conditions. The primers for genotyping were F-TGTCCAACAGAACAGTGAACTGGAG and R-CCAAGCTCTTTAAGCCGACAGTG. All animal experiments were conducted in accordance with the Regulations of Hubei Province Laboratory Animal Management and approved by Wuhan University Animal Experiment Ethics Committee.

### Preparation of bone marrow-derived dendritic cells (BMDC)

Bone marrow cells were isolated from C57BL/6J mouse tibia and femur and then cultured for 7–9 days in 10% FBS DMEM containing mouse GM-CSF (50 ng/mL, Peprotech).

### Preparation of bronchoalveolar lavage fluid (BALF) and RNA-seq library construction and sequencing

The methods were the same as previously described (39). NSUN2 expression analysis in COVID-19 patients compared with healthy individuals was obtained from the analysis of previous results (https://github.com/zhouyulab/ncov/).

### Plasmids and RNA interference

NSUN2 was cloned into both the pCAGGS and pGEX6P-1 vector. The sequences of siRNAs were si-h-NSUN2#1: GAGATCCTCTTCTATGATC; si-h-NSUN2#2: GGAGAACAAGCTGTTCGAG; si-h-TRDMT1: GCGATATGCTCTTCT GTTA; si-h-METTL3: CTGCAAGTATGTTCACTATGA; si-h-METTL14: AAGGATGAGTTAATAGCTAAA; si-h-ALKBH5: GTCGGGACTGCATAATTAA. Cells were seeded and siRNAs were transfected using Pepmute. The knockdown efficiency was detected 36 h after transfection using immunoblot analysis or qPCR.

### Antibodies and immunoblot analysis

The antibodies used were as follows: rabbit anti-NSUN2 (Proteintech, 20854-1-AP), rabbit anti-Phospho-IRF-3-Ser396 (CST, 83611S), rabbit anti-IRF3 (Proteintech, 11312-1-AP), rabbit anti-phospho-TBK1/NAK-Ser172 (CST, 14590S), rabbit anti-TBK1/NAK (CST, 38066S), mouse anti-HA (Sigma, H6908), rabbit anti-HA (Sigma, H3663), mouse anti-Flag (Proteintech, 66008-3-Ig), rabbit anti-Flag (Sigma, SAB4301135), mouse anti-m^5^C antibody (Abcam, ab10805), mouse anti-GAPDH (Proteintech, 60004-1-Ig), mouse anti-β-actin (Proteintech, 66009-1-Ig). Cells were washed once with PBS and lysed in RIPA lysis buffer (50 mM Tris, pH 7.6, 1% NP-40, 150 mM NaCl, 0.1% SDS). 5× SDS loading buffer was added to the protein sample and boiled for 5 min. Samples were resolved on SDS-PAGE and transferred onto nitrocellulose membrane (GE Healthcare), followed by blocking with TBS containing 0.1% Tween-20 (TBST) and 5% non-fat powdered milk or bovine serum albumin (BSA) and probing with different antibodies.

### Co-immunoprecipitation and RNA-binding protein immunoprecipitation (RIP)

HEK293T cells were seeded onto 6 cm dishes and transfected as illustrated above. Thirty-six hours after transfection, cells were lysed in RIPA buffer (50 mM Tris, pH 7.6, 1% NP-40, 150 mM NaCl, 0.1% SDS) containing protease inhibitors and phosphatase inhibitors, if necessary. The cell lysates were incubated overnight at 4 °C with HA-tag rabbit mAb beads (Sepharose Bead Conjugate, 3956S, CST) or Flag-tag rabbit mAb beads (Sepharose Bead Conjugate, 70569S, CST). The beads were washed five times with cold PBS and then mixed with SDS loading buffer and boiled for 10 min prior to SDS-PAGE and immunoblot analysis. For RNA immunoprecipitation, HEK293T cells were transfected and lysed with lysis buffer (20 mM Tris–HCl, pH 7.6, 150 mM NaCl, 1% Triton-X100, 1 mM EDTA, 0.1% SDS, and 2 mM DTT, RNase free) and incubated overnight at 4 °C with HA-tag rabbit mAb beads. Beads were washed five times with lysis buffer and divided in half for RNA extraction and qPCR analysis or for immunoblot analysis.

### RNA isolation and qPCR

Total RNA was isolated using TRIzol reagent (Invitrogen) following the manufacturer’s instructions. The isolated mRNA was reverse transcribed to cDNA using PrimeScript RT Reagent Kit (Takara, RR037A). Real-time quantitative PCR was carried out through ABI 7500 Real Time PCR System by SYBR Green Master Mix (YEASEN, 11199ES03). GAPDH was used in normalization via the ΔΔCt method. Primer sequences are shown in Supplementary Table S1.

### Protein expression and purification

*Escherichia coli* BL21 cells were transformed with pGEX-6p-1-GST-NSUN2 and cultured in 10 mL Luria broth medium at 37 °C for 6 h. The culture was then transferred to 1000 mL Luria broth medium and grown at 37 °C to an absorbance of 0.6–1 as measured at 600 nm. IPTG was added to the culture to achieve a final concentration of 0.2 mM and induced at 16 °C for 16–20 hours. Cell cultures were harvested by centrifugation and then lysed by lysozyme and ultrasonication. GST-tagged NSUN2 proteins were purified by affinity chromatography using reduced glutathione resin (GenScript, L00206) following the manufacturer’s instructions. Finally, the recombinant proteins were eluted through incubation for 30 min at 4 °C with 100 μL of 50 mM Tris (pH 8.0), 2 mM DTT and 10 mM reduced glutathione and 8% glycerine was added for snap-freezing in liquid nitrogen and storage at −80 °C until use. The purity and quantity of the recombinant proteins were assessed by SDS-PAGE followed by staining with Coomassie blue and immunoblot analysis.

### *In vitro* transcription assays

The cDNA of HeLa cells was used as a template for PCR amplification of each segment of IRF3, which were then used as templates for *in vitro* transcription following the manufacturer’s instructions (Invitrogen, 00612295). All 5’ primers of the segments contained the T7 promoter sequence (TAATACGACTCACTATAGGG). The transcription reaction was performed at 30 °C for 16 h. The transcribed RNA was precipitated and identified by agarose gel electrophoresis.

### *In vitro* methylation assays

Reaction mixtures (50 µL) containing 0.2 nM recombinant GST-tagged NSUN2, 0.01 nM *in vitro* transcribed fragments of mRNA, 1 µCi of *S*-adenosyl [methyl-^3^H] methionine (0.5 μCi/μl; PerkinElmer) in reaction buffer (500 mM Tris–HCl (pH 7.5), 5 mM EDTA, 10 mM dithiothreitol, 20 mM MgCl_2_) and 40 units of RNase inhibitor were incubated for 60 min at 37 °C, as described (20). The ^3^H-labeled products were isolated using DEAE-Sephadex A-50 columns and quantitated by liquid scintillation counting (PerkinElmer). Non-isotopic methylated RNA fragments were prepared using cold SAM (Biolabs, 0991410) and *in vitro* transcribed RNA fragments under similar conditions.

### Reporter gene assays

Cells were seeded into 24-well plates (2 × 10^5^ cells per well) and transfected with 100 ng of luciferase reporter plasmid together with a total of 0.5 μg of expression plasmid or empty control plasmid via Lipofectamine 2000 or Neofect. Twenty nanograms of pRL-TK *Renilla* luciferase reporter plasmid was also transfected to normalize the transfection efficiency. For the knockdown system, siRNAs were first transfected by Pepmute, and 24 hours later, luciferase reporter plasmid and pRL-TK *Renilla* luciferase reporter were subsequently transfected by Lipofectamine 2000. Luciferase activity in total cell lysates was measured using a dual-luciferase reporter assay system (Promega).

### VSV plaque assay

Vero cells were seeded into 24-well plates to about 80%–90% density before infection. The supernatants containing VSV then were serially diluted for infection of Vero cells. Two hours later, supernatants were removed, and PBS was used to wash the infected Vero cells. The DMEM containing 2% methylcellulose and 10% FBS was overlaid onto the cells. Two days later, cells were fixed and stained with formaldehyde (4%) and crystal violetin (0.2%) for 6 h followed by washing with water. Finally, plaques were counted, and the results were averaged and multiplied by the dilution factor for calculation of viral titers as PFU/mL and statistical analyses were performed.

### Endogenous *IRF3* mRNA pull down

The four IRF3 CHIRP probes were as follows: CTTTATCATTCTTTGGGTAACA, AACTCGTAGATTTTATGTGGGT, AGATGGTCTGCTGGAAGACTTG, and AGGAACCAGTTTATTGGTTGAG. All the probes were 3’biotin-TEG-modified (Sangon company). Ten × 10 cm dishes of cells were used for total RNA extraction for each group. The total RNA was dissolved in 600 µL hybrid buffer (350 mM NaCl, 0.5% SDS, 25 mM Tris–HCl, 1 mM EDTA, 7.5% formamide, pH 7.5), and 5 µL IRF3 probes (100 μM) were added and incubated at 65 ℃ for 5 min followed by 37 ℃ while rotating for 2 hours. Then, 100 µL Dynabeads M-280 streptavidin (ThermoFisher, 11205D) was added followed by rotating at 37 ℃ for 1 h. Six hundred microliters of wash buffer (2× SSC buffer, 0.5% SDS, RNase inhibitor) was used to wash the beads 5 times for 5 min at 4 ℃. RNase-free water (20 μL) was added for elution followed by incubation at 75 ℃ for 5 min. After centrifuging at 1000*g* for 3 min, the pulled down RNA was got in the eluate supernatant.

### m^5^C Dot blot analysis

Equal amounts mRNA were denatured at 65 °C for 10 min followed by immediate chilling on ice. mRNA was mixed with RNA loading buffer and then carefully spotted onto a Hybond-N+ membrane (GE Healthcare), followed by UV crosslinking. The membranes were washed with TBST 2 times and blocked with 5% BSA in TBST for 2 hours. The anti-m^5^C antibody (Abcam, ab10805) was diluted 1:500 and incubated with the membranes at 4 °C overnight. Membranes were washed 3 times with TBST for 10 min and then incubated with goat anti-mouse IgG-HRP for 1 hour at room temperature. Membranes were washed 3 times with TBST for 5 min followed by chemiluminescence. Equal RNA loading was verified by methylene blue (MB) staining.

### m^5^C-Methylated RNA immunoprecipitation (MeRIP)

For MeRIP, 200 μg of total RNA was incubated with anti-m^5^C antibody in 800 µL of IPP buffer (150 mM NaCl, 0.1% NP-40, 10 mM Tris–HCl, pH 7.4) for 2 h at 4 °C. The mixture was then incubated with 30 µL proteinA/G beads overnight. The beads were then washed 5 times with IPP buffer, followed by RNA extraction and qPCR analysis.

### Bisulfite RNA sequencing

The adaptor sequences used were Adaptor-F: AGGTCTGGCTGAAGTTGA; Adaptor-R: ATACCTCCGTGACCATTT. The sequencing primers were Adaptor-F-mut: AGGTTTGGTTGAAGTTGA; Adaptor-R-mut: ATACCTCCATAACCATTT. Bisulfite RNA sequencing was performed to identify the m^5^C methylation site within an RNA fragment as previously described (48, 49). Briefly, 10 μg in vitro methylated RNA fragment (methylated by NSUN2 using cold SAM or unmethylated) was dissolved in 10 µL of RNase-free water and denatured at 65 °C for 10 min followed by immediate chilling on ice. Samples were then mixed with 42.5 µL of 5 M sodium bisulfite mix (Epitect) and 17.5 µL DNA protection buffer (Epitect) and incubated at 70 °C for 5 min then 60 °C for 1 hour, and this process was repeated for 4 cycles, followed by desalting using Micro Bio-spin 6 Chromatography Columns (Biorad, 732-6200). Then, the RNA adducts were desulfonated by adding 1 volume of Tris–HCl (pH 9.0) at 37 °C for 1 h. Next, 0.3 M sodium acetate (pH 5.2), 20 μg glycogen (Beyotime, D0812) and 3 volumes of 100% ethanol were added for precipitation. The RNA was precipitated at −80 °C for at least 5 h and then centrifuged. The bisulfite-converted RNA was reverse-transcribed using Adaptor-R-mut primer and random primer and subjected to PCR with Es Taq DNA polymerase (CW0688S) using Adaptor-mut primer pairs. The PCR products were inserted into the pGEM-T Easy Vector System (Promega, A1360) following the manufacturer’s instructions. The plasmids purified from single clones were sequenced by T7 promoter. The sequencing results were checked by alignment with the corresponding original *IRF3* mRNA sequence, and the retained cytosines (C) were considered to be methylated by NSUN2. The unmethylated cytosines (C) were converted to uracils (U) on RNA segments.

### Lentiviral Package and Infection

A lentiviral system was utilized to obtain NSUN2 knockout cells or stable cell lines in *Irf3^−/−^Irf7^−/−^* MEFs. For this, lentiviral backbone (2 µg), psPAX2 (1 μg), and pMD2.G (1 μg) were transiently transfected into HEK293T cells which were plated on 6-well plates. Forty-eight hours later, supernatants were collected and filtered using a 0.45 μm filter to infect target cells with polybrene (8 µg/mL). Cells were infected twice to get a higher transduction efficiency. Then, puromycin was used to screen positive cells.

### Construction of knockout cell line by CRISPR/Cas9

The gRNAs were NSUN2-gRNA-1: F-CACCGACGCGGAGGATGGCGCCGA and R-AAACTCGGCGCCATCCTCCGCGTC; NSUN2-gRNA-2: F-CACCACCGTG GCGTTTCAGCGGTT and R-AAACAACCGCTGAAACGCCACGGT. The gRNAs were constructed in lentiCRISPR-v2 plasmid (Addgene). The lentiviral package and infection were the same as above, followed by seeding into 96-well plates (1 cell per well). After two weeks’ cultivation, single clones were selected following enlarged cultivation with puromycin selection. Single clones were identified by immunoblot analysis, and genomic DNA was extracted followed by PCR and sequencing.

### Ethics statement

This study was approved by the Ethics Committee of the Zhongnan Hospital of Wuhan University. The RNA-seq analyses of BALF samples were performed on existing samples collected during standard diagnostic tests, posing no extra burden to patients.

## Acknowledgement

We thank Yingle Liu and Mang Shi for providing BALF samples of COVID-19 patients. We thank Dr. Zheng-Li Shi for providing SARS-CoV-2, Dr. Hong-Bing Shu for providing SeV, HSV-1, VSV-GFP, Dr. Bo Zhang for providing ZIKV, and Dr. Ming-Zhou Chen for providing VSV. This study was supported by grants from the National Science and Technology Major Project (2018YFA0900801), China NSFC projects (32041007 and 81672008), Hubei Natural Science Foundation (2018CFA035), Basic Scientific Research Foundation of Central Universities (2042019gf0026) and Special Fund for COVID-19 Research of Wuhan University. We are grateful to Beijing Taikang Yicai Foundation for their great support to this work.

## Author contributions

Y.C. and H.W. conceived the research and experiments. H.W., C.Z., L.Z., J.F. and M.H. performed the major experiments and analysis. Y.Z., K.L. and D.W. analyzed transcriptome sequencing data. C.F., H.T. and A.J. provided critical advice. H.W. and Y.C. wrote the manuscript with contributions from all other authors.

## Supplementary information

**Supplementary Fig S1.**
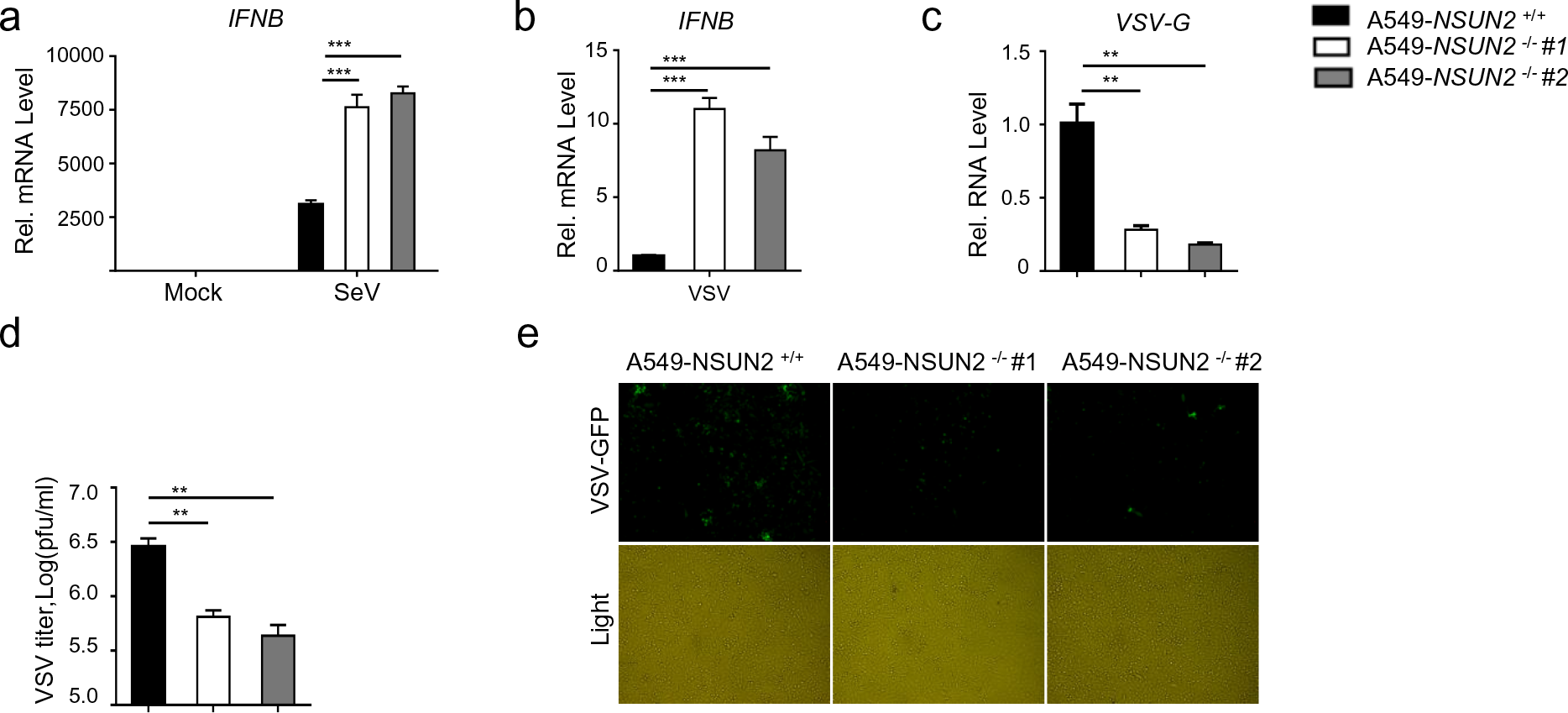
Knockout of NSUN2 in A549 cells promotes IFN-β responses. (a) qPCR analysis of *IFNB* mRNA in wild-type A549 cells or NSUN2 knockout A549 cells, with or without infection by SeV, for 8 h. (b-c) qPCR analysis of *IFNB* mRNA or *VSV-G* RNA in wild-type A549 cells or NSUN2 knockout A549 cells, with infection by VSV for 24 h (MOI = 0.005). (d) VSV plaque assay in in wild-type A549 cells or NSUN2 knockout A549 cells, with infection by VSV for 24 h (MOI = 0.005). (e) Microscopy analysis of VSV-GFP replication in wild-type A549 cells or *NSUN2^−/−^* A549 cells, all infected with VSV-GFP for 18 h (MOI = 0.005). Data are representative of three independent experiments and analyzed by two-tailed unpaired t test. Graphs show the mean ± SD (n = 3) derived from three independent experiments. NS, not significant for *P* > 0.05, \**P* < 0.05, \*\**P* < 0.01, \*\*\**P* < 0.001.

**Supplementary Fig S2.**
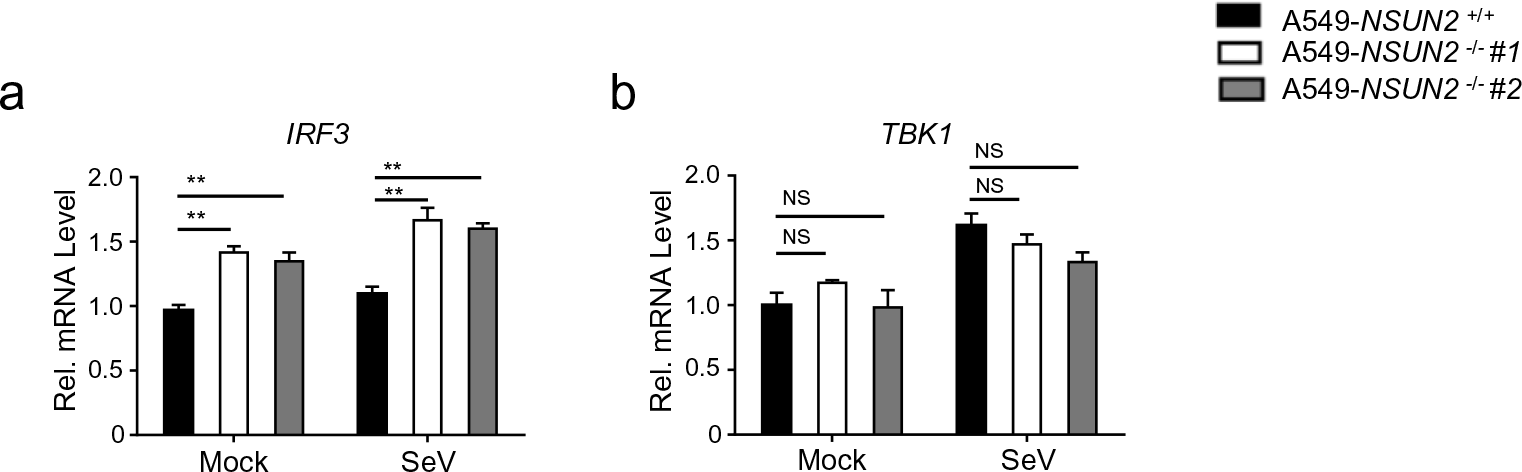
Knockout of NSUN2 in A549 cells elevated the endogenous *IRF3* mRNA level. (a-b) qPCR analysis of *IRF3* or *TBK1* mRNA in wild-type A549 cells or NSUN2 knockout A549 cells, with or without infection by SeV, for 8 h.

**Supplementary Fig S3.**
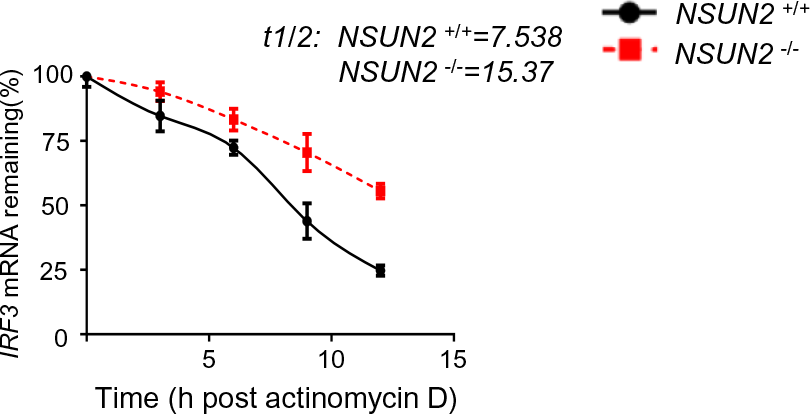
Knockout of NSUN2 enhances *IRF3* mRNA stability in A549 cells. (a) Stability analysis of *IRF3* mRNA in wild-type A549 cells or *NSUN2^−/−^* A549 cells with treatment of actinomycin D (ActD) for 0, 3, 6, 9, and 12 h. Data are representative of three independent experiments and were analyzed by two-tailed unpaired t test. Graphs show the mean ± SD (n = 3) derived from three independent experiments. NS, not significant for *P* > 0.05, \**P* < 0.05, \*\**P* < 0.01, \*\*\**P* < 0.001.

**Supplementary Fig S4.**
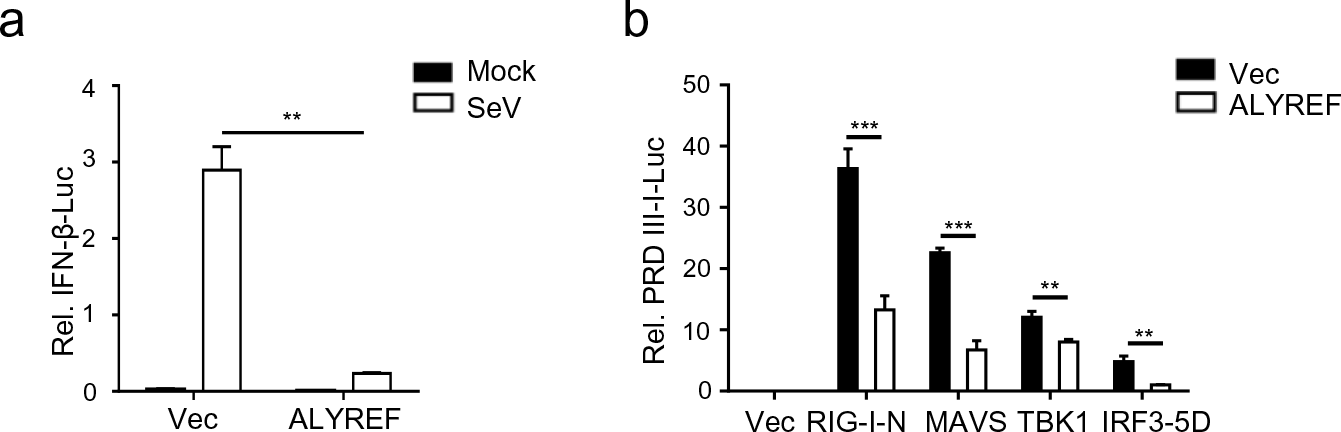
ALYREF negatively regulates IFN-β response. (a) Dual-luciferase analysis of IFN-β-Luc activity in HEK293T cells in 24-well plates transfected for 24 h with 100 ng IFN-β firefly luciferase reporter (IFN-β-Luc) and 20 ng *Renilla* luciferase (RL-TK), along with vector or the plasmid encoding ALYREF, with or without infection by SeV, for another 10 h. (b) Dual-luciferase analysis of PRDIII-I-Luc activity in HEK293T cells in 24-well plates transfected for 24 h with the indicated RIG-N, MAVS, TBK1, and IRF3-5D expression plasmids with co-transfection with vector or ALYREF. Data are representative of three independent experiments and were analyzed by two-tailed unpaired t test. Graphs show the mean ± SD (n = 3) derived from three independent experiments. NS, not significant for *P* > 0.05, \**P* < 0.05, \*\**P* < 0.01, \*\*\**P* < 0.001.

**Supplementary Fig S5.**
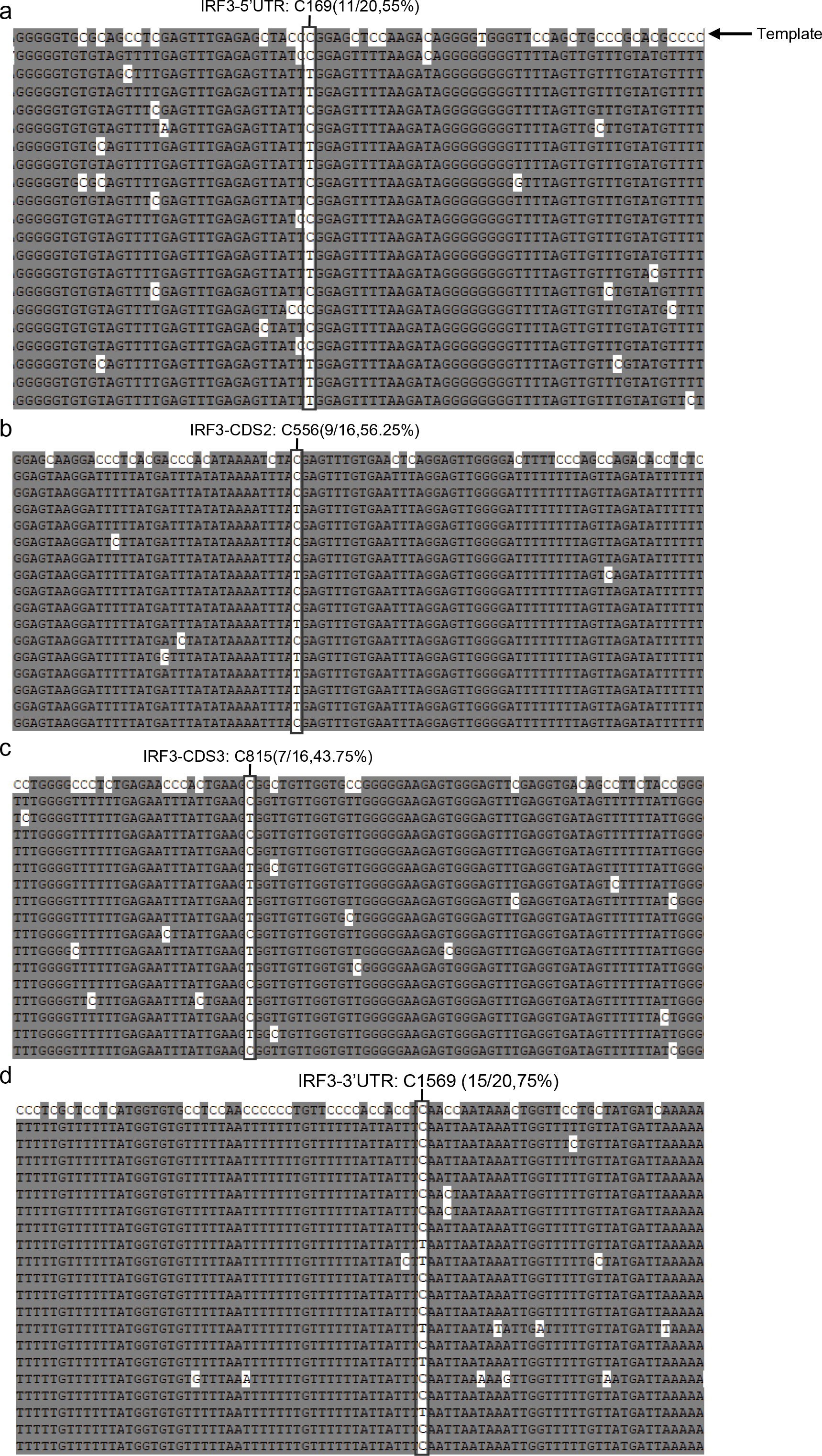
Bisulfite sequencing alignment of IRF3 segments. (a-d) The *in vitro*transcribed IRF3 segments were subjected to NSUN2 methylation and bisulfite RNA sequencing. The sequence on the top was the original template, and the lower sequences were the identified segments. The retained cytosines (C) were considered to be methylated by NSUN2, while the unmethylated cytosines (C) were converted to uracils (U) which then converted to thymines (T) after PCR. The ratio of m^5^C to (C + m^5^C) represents the methylation rate.

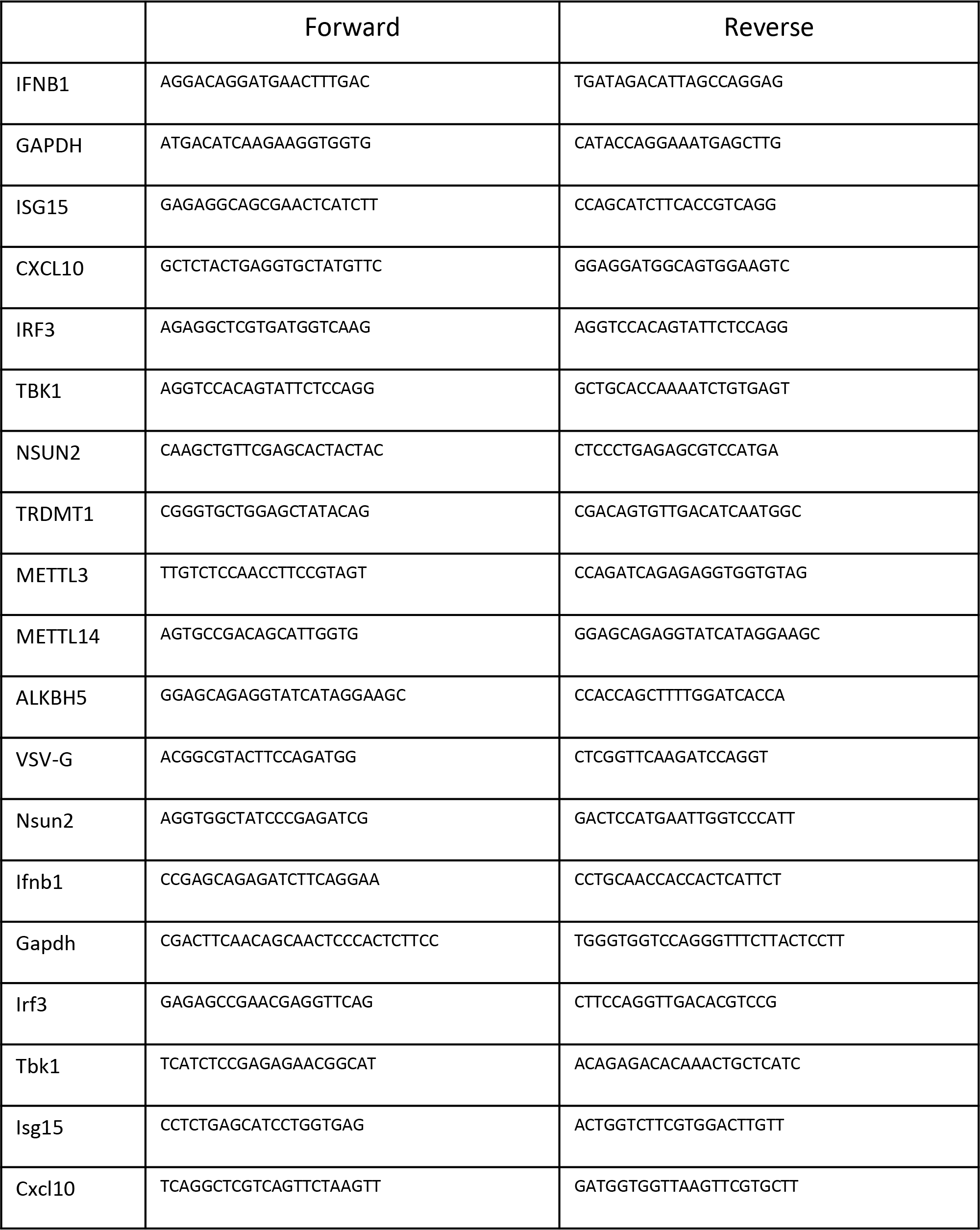
Primers for mRNA Quantification

## References

1. Roundtree IA, Evans ME, Pan T, & He C (2017) Dynamic RNA Modifications in Gene Expression Regulation. Cell 169(7):1187–1200.

2. Dezi V, Ivanov C, Haussmann IU, & Soller M (2016) Nucleotide modifications in messenger RNA and their role in development and disease. Biochem Soc Trans 44(5):1385–1393.

3. Trixl L & Lusser A (2019) The dynamic RNA modification 5-methylcytosine and its emerging role as an epitranscriptomic mark. Wiley Interdiscip Rev RNA 10(1):e1510.

4. Chellamuthu A & Gray SG (2020) The RNA Methyltransferase NSUN2 and Its Potential Roles in Cancer. Cells 9(8).

5. Yang Y, Hsu PJ, Chen YS, & Yang YG (2018) Dynamic transcriptomic m(6)A decoration: writers, erasers, readers and functions in RNA metabolism. Cell Res 28(6):616–624.

6. Cao G, Li HB, Yin Z, & Flavell RA (2016) Recent advances in dynamic m6A RNA modification. Open Biol 6(4):160003.

7. Yang J, Wang H, & Zhang W (2019) Regulation of Virus Replication and T Cell Homeostasis by N(6)-Methyladenosine. Virol Sin 34(1):22–29.

8. Han D, et al. (2019) Anti-tumour immunity controlled through mRNA m(6)A methylation and YTHDF1 in dendritic cells. Nature 566(7743):270–274.

9. Frye M, Harada BT, Behm M, & He C (2018) RNA modifications modulate gene expression during development. Science 361(6409):1346–1349.

10. Winkler R, et al. (2019) m6A modification controls the innate immune response to infection by targeting type I interferons (vol 20, pg 173, 2018). Nature Immunology 20(2):243–243.

11. Rubio RM, Depledge DP, Bianco C, Thompson L, & Mohr I (2018) RNA m(6) A modification enzymes shape innate responses to DNA by regulating interferon beta. Genes Dev 32(23-24):1472–1484.

12. Liu Y, et al. (2019) N6-methyladenosine RNA modification–mediated cellular metabolism rewiring inhibits viral replication. Science 365(6458):1171–1176.

13. Zheng Q, Hou J, Zhou Y, Li Z, & Cao X (2017) The RNA helicase DDX46 inhibits innate immunity by entrapping m(6)A-demethylated antiviral transcripts in the nucleus. Nat Immunol 18(10):1094–1103.

14. Wang L, Wen M, & Cao X (2019) Nuclear hnRNPA2B1 initiates and amplifies the innate immune response to DNA viruses. Science 365(6454):eaav0758.

15. Squires JE, et al. (2012) Widespread occurrence of 5-methylcytosine in human coding and non-coding RNA. Nucleic Acids Res 40(11):5023–5033.

16. Tuorto F, et al. (2012) RNA cytosine methylation by Dnmt2 and NSun2 promotes tRNA stability and protein synthesis. Nat Struct Mol Biol 19(9):900–905.

17. Zhang XT, et al. (2012) The tRNA methyltransferase NSun2 stabilizes p16(INK4) mRNA by methylating the 3’-untranslated region of p16. Nature Communications 3.

18. Mei L, et al. (2020) RNA methyltransferase NSUN2 promotes gastric cancer cell proliferation by repressing p57(Kip2) by an m(5)C-dependent manner. Cell Death Dis 11(4):270.

19. Tang H, et al. (2015) NSun2 delays replicative senescence by repressing p27 (KIP1) translation and elevating CDK1 translation. Aging (Albany NY) 7(12):1143–1158.

20. Li Q, et al. (2017) NSUN2-Mediated m5C Methylation and METTL3/METTL14-Mediated m6A Methylation Cooperatively Enhance p21 Translation. J Cell Biochem 118(9):2587–2598.

21. Schumann U, et al. (2020) Multiple links between 5-methylcytosine content of mRNA and translation. BMC Biol 18(1):40.

22. Yang X, et al. (2017) 5-methylcytosine promotes mRNA export-NSUN2 as the methyltransferase and ALYREF as an m(5)C reader. Cell Research 27(5):606–625.

23. Chen X, et al. (2019) 5-methylcytosine promotes pathogenesis of bladder cancer through stabilizing mRNAs. Nat Cell Biol 21(8):978–990.

24. Yang Y, et al. (2019) RNA 5-Methylcytosine Facilitates the Maternal-to-Zygotic Transition by Preventing Maternal mRNA Decay. Mol Cell 75(6):1188–1202 e1111.

25. Zou F, et al. (2020) Drosophila YBX1 homolog YPS promotes ovarian germ line stem cell development by preferentially recognizing 5-methylcytosine RNAs. Proc Natl Acad Sci U S A 117(7):3603–3609.

26. Eckwahl M, et al. (2020) 5-Methylcytosine RNA Modifications Promote Retrovirus Replication in an ALYREF Reader Protein-Dependent Manner. J Virol 94(13).

27. Honda K, Takaoka A, & Taniguchi T (2006) Type I interferon [corrected] gene induction by the interferon regulatory factor family of transcription factors. Immunity 25(3):349–360.

28. Ablasser A & Hur S (2019) Regulation of cGAS- and RLR-mediated immunity to nucleic acids. Nature Immunology 21(1):17–29.

29. Fitzgerald KA & Kagan JC (2020) Toll-like Receptors and the Control of Immunity. Cell 180(6):1044–1066.

30. Rehwinkel J & Gack MU (2020) RIG-I-like receptors: their regulation and roles in RNA sensing. Nat Rev Immunol 20(9):537–551.

31. Honda K & Taniguchi T (2006) IRFs: master regulators of signalling by Toll-like receptors and cytosolic pattern-recognition receptors. Nat Rev Immunol 6(9):644–658.

32. Wu J & Chen ZJ (2014) Innate immune sensing and signaling of cytosolic nucleic acids. Annu Rev Immunol 32:461–488.

33. Tamura T, Yanai H, Savitsky D, & Taniguchi T (2008) The IRF family transcription factors in immunity and oncogenesis. Annu Rev Immunol 26:535–584.

34. Schneider WM, Chevillotte MD, & Rice CM (2014) Interferon-stimulated genes: a complex web of host defenses. Annu Rev Immunol 32:513–545.

35. Li S, et al. (2016) The tumor suppressor PTEN has a critical role in antiviral innate immunity. Nat Immunol 17(3):241–249.

36. Mancino A & Natoli G (2016) Specificity and Function of IRF Family Transcription Factors: Insights from Genomics. J Interferon Cytokine Res 36(7):462–469.

37. Cao Y, et al. (2018) PTEN-L promotes type I interferon responses and antiviral immunity. Cell Mol Immunol 15(1):48–57.

38. Zhou Y, et al. (2019) Interferon-inducible cytoplasmic lncLrrc55-AS promotes antiviral innate responses by strengthening IRF3 phosphorylation. Cell Res 29(8):641–654.

39. Xiong Y, et al. (2020) Transcriptomic characteristics of bronchoalveolar lavage fluid and peripheral blood mononuclear cells in COVID-19 patients. Emerg Microbes Infect 9(1):761–770.

40. Yuan S, et al. (2014) Methylation by NSun2 represses the levels and function of microRNA 125b. Molecular and cellular biology 34(19):3630–3641.

41. Huai W, et al. (2019) KAT8 selectively inhibits antiviral immunity by acetylating IRF3. J Exp Med 216(4):772–785.

42. Wang P, Zhao W, Zhao K, Zhang L, & Gao C (2015) TRIM26 negatively regulates interferon-beta production and antiviral response through polyubiquitination and degradation of nuclear IRF3. PLoS Pathog 11(3):e1004726.

43. Moon HJ & Redman KL (2014) Trm4 and Nsun2 RNA:m5C methyltransferases form metabolite-dependent, covalent adducts with previously methylated RNA. Biochemistry 53(45):7132–7144.

44. Wang X, et al. (2014) N6-methyladenosine-dependent regulation of messenger RNA stability. Nature 505(7481):117–120.

45. Shi H, et al. (2017) YTHDF3 facilitates translation and decay of N(6)-methyladenosine-modified RNA. Cell Res 27(3):315–328.

46. Wang X, et al. (2015) N(6)-methyladenosine Modulates Messenger RNA Translation Efficiency. Cell 161(6):1388–1399.

47. Xiao W, et al. (2016) Nuclear m(6)A Reader YTHDC1 Regulates mRNA Splicing. Mol Cell 61(4):507–519.

48. Pollex T, Hanna K, & Schaefer M (2010) Detection of cytosine methylation in RNA using bisulfite sequencing. Cold Spring Harb Protoc 2010(10):pdbprot5505.

49. Schaefer M, Pollex T, Hanna K, & Lyko F (2009) RNA cytosine methylation analysis by bisulfite sequencing. Nucleic Acids Res 37(2):e12.

